# Single-Cell Epitope-Transcriptomics Reveal Lung Stromal and Immune Cell Response Kinetics to Nanoparticle-delivered RIG-I and TLR4 Agonists

**DOI:** 10.1101/2022.11.29.518316

**Authors:** M. Cole Keenum, Paramita Chatterjee, Alexandra Atalis, Bhawana Pandey, Angela Jimenez, Krishnendu Roy

**Affiliations:** Wallace H. Coulter Department of Biomedical Engineering, Georgia Institute of Technology and Emory University, Atlanta, Georgia, USA; Marcus Center for Therapeutic Cell Characterization and Manufacturing, Georgia Institute of Technology, Atlanta, Georgia, USA; The Parker H. Petit Institute for Bioengineering and Biosciences, Georgia Institute of Technology, Atlanta, Georgia, USA

**Keywords:** RIG-I lung, TLR4 lung, innate immune response, lung immune response, single cell lung, interferon signaling lung, ribosome biogenesis immune, ion immune response

## Abstract

Lung-resident and circulatory lymphoid, myeloid, and stromal cells, expressing various pattern recognition receptors (PRRs), detect pathogen and danger-associated molecular patterns (PAMPs/DAMPs), and defend against respiratory pathogens and injuries. Here, we report the early responses of murine lungs to nanoparticle-delivered PAMPs, specifically the RIG-I agonist poly-U/UC (PUUC), with or without the TLR4 agonist monophosphoryl lipid A (MPLA). Using cellular indexing of transcriptomes and epitopes by sequencing (CITE-seq), we characterized the responses at 4 and 24 hours after intranasal administration. Within 4 hours, ribosome-associated transcripts decreased in both stromal and immune cells, followed by widespread interferon-stimulated gene (ISG) expression. Using RNA velocity, we show that lung-neutrophils dynamically regulate the synthesis of cytokines like CXCL-10, IL-1α, and IL-1β. Co-delivery of MPLA and PUUC increased chemokine synthesis and upregulated antimicrobial binding proteins targeting iron, manganese, and zinc in many cell types, including fibroblasts, endothelial cells, and epithelial cells. Overall, our results elucidate the early PAMP-induced cellular responses in the lung and demonstrate that stimulation of the RIG-I pathway, with or without TLR4 agonists, induces a ubiquitous microbial defense state in lung stromal and immune cells. Nanoparticle-delivered combination PAMPs may have applications in intranasal antiviral and antimicrobial therapies and prophylaxis.

## INTRODUCTION

Respiratory infections are among the leading causes of death and illness worldwide.^1,2^ Because the respiratory system handles thousands of liters of nonsterile air daily, it is equipped with a coordinated, fast-acting immune system to respond to pathogens.^3^ Whereas many studies typically focus on traditional immune-cell populations like granulocytes, monocytes, and lymphocytes, increasing evidence shows that structural cells from the endothelium, epithelium, and stroma are primed for immune functions, including those in the lungs.^4^

In the lower respiratory tree patrolling alveolar macrophages (AMs) phagocytose pathogens that survive upper airway filtration.^5^ AMs roam over the epithelium, where flat alveolar type I (AT1) pneumocytes cover most of the thin epithelial barrier between pathogens and the lung vasculature. Alveolar type II (AT2) pneumocytes, which are smaller yet more numerous than AT1 cells,^6^ are bipotent epithelial progenitors that secrete surfactant containing proteins A and D. These proteins are soluble pattern recognition receptors (PRRs) that contribute to antimicrobial responses by opsonizing microbes and interacting with dendritic cells (DCs), macrophages, T cells, and neutrophils.^7-9^ On the interstitial side of the alveoli are aerocyte (aCap), flattened endothelial cells specialized for gas exchange and leukocyte transport, and general capillary endothelial cells (gCap), bipotent progenitors that may present antigen via MHC class II.^10^ Between the epithelial and endothelial layers, the interstitial macrophages provide a regulatory function and may suppress the development of allergic pathologies like asthma.^11^ Also in the interstitium, plasmacytoid and other DCs express high levels of PRRs that sense microbial PAMPs and further activate other immune cells such as cytotoxic and helper T cells, natural killer (NK) cells, and natural killer T (NKT) cells.^3^

Importantly, many other lung cells express PRRs. Structural cells like alveolar epithelial cells and lung fibroblasts express TLR4. They detect gram-negative bacterial lipopolysaccharide (LPS), leading to NF-κB activation and synthesis of proinflammatory cytokines and type I interferon.^12-14^ TLR4 from the endothelium, not leukocytes, is the primary driver of neutrophil sequestration in the LPS-stimulated lung.^15^ Intracellularly, the PRR class of retinoic acid-inducible gene I (RIG-I)-like receptors (RLRs) sense foreign nucleic acids to initiate antiviral immunity.^16^ All cell types express RLRs,^17^ which serve to recognize RNA viruses, including SARS-CoV-2.^18,19^ RIG-I signaling converges on the activation of transcription factors IRF-3, IRF-7, and NF-kB to produce type I interferons. Once secreted, these interferons ultimately drive the synthesis of interferon-stimulate genes (ISGs) to create an antiviral state.^16^ This antiviral state classically includes the transcription of genes, including *Eif2ak2* (protein kinase RNA-activated, PKR), which phosphorylates and inhibits ribosomal eIF2-α, and members of the *Oas1* family (2′,5′-oligoadenylate synthetase, OAS), which activates RNase L to degrade cellular mRNA globally.^20,21^ These are some of the mechanisms by which ISGs post-translationally inhibit host and viral mRNA translation. However, whether ribosomes are regulated transcriptionally in an antiviral state is poorly understood.

Lung PRR ligation also regulates antimicrobial responses by upregulating genes related to metal ion homeostasis. For example, at baseline, alveolar epithelial cells and AMs express isoforms of divalent metal ion transporter 1 (DMT1) and ferroportin 1 (FPN1), which regulate iron transport. LPS induces hepcidin expression in AMs, which sequesters iron and prevents its usage by invading microbes.^22^ In acute immune responses, alveolar epithelial cells express *Lcn2* (lipocalin-2/siderocalin) which binds to bacterial siderophores and contributes to bacteriostatic responses against airway pathogens like *Mycobacterium tuberculosis* and *Klebsiella pneumoniae*.^23,24^

To study innate immune responses to PRR ligation by PAMPs in the mouse lung, we intranasally administered the RIG-I agonist PUUC with and without the TLR4 agonist MPLA using poly(lactic-co-glycolic acid)-polyethyleneimine-based nanoparticles (PLGA-PEI NPs). We have previously reported that these PLGA-PEI pathogen-like particles (PLPs) can simultaneously activate multiple PRRs,^25,26^ including intracellular RIG-I and extracellular TLR4 *in vitro* and *in vivo*.^*27*^ After intranasal administration with PUUC NPs or MPLA+PUUC NPs, we employed PCR microarray analysis to survey the general transcriptional landscape of the lung at 4 and 24 h. We then used Cellular Indexing of Transcriptomes and Epitopes (CITE-Seq) to define cell types according to surface protein and mRNA expression,^28^ perform differentially expressed gene (DEG) analysis, gene set enrichment analysis (GSEA), RNA velocity analysis of neutrophils, and cell-cell signaling analysis at the same time points. Analyses were conducted comparing each treatment against naïve lungs and for the MPLA+PUUC-treated lungs against PUUC-treated lungs. Our investigation targeted essentially all identified cell types in the lung, including epithelial cells, fibroblasts, endothelial cells, granulocytes, monocytes, and lymphocytes. We uncovered both cell-type specific and general immune functions in the lung at these early time points, including regulation of ribosome-associated genes, variable expression of proinflammatory cytokines by structural cells, and regulation of ion- and metal-binding proteins especially in the context of MPLA+PUUC treatment. Overall, we found that particulate delivery of PUUC alone or MPLA and PUUC together can produce a broadly antimicrobial state with transcriptional responses starting within hours in all cell types identified. Delivery of such particles, containing PAMPs that can be mixed and combined, could be further investigated as a prophylactic treatment to generate a synthetic antiviral or antibacterial state. Or, since microbes commonly downregulate the pathways that would otherwise be upregulated by PAMP ligation of PRRs, these particles could be investigated for uses as a pan-antiviral or pan-antibacterial therapy.

## METHODS

### PUUC and MPLA+PUUC PLP Synthesis

PLGA-PEI NPs were synthesized as previously described.^27^ Briefly, PLGA (50 lactate:50 glycolate, MW: 7000–17,000, Resomer RG 502H, Sigma-Aldrich, #719897) was dissolved in dichloromethane (DCM, Sigma-Aldrich, #270997) and endotoxin-free water in a 1:20:5 w/v/v ratio to form a primary emulsion that was sonicated at 65% power for 2 min. For particles with encapsulated TLR4 agonist, MPLA PHAD® (1 μg/mg PLGA, Avanti Polar Lipids, #699800P) was dissolved in DCM prior to emulsification. This first emulsion was added to 5% polyvinyl alcohol (M_w_ 31,000-50,000, 87-89% hydrolyzed, SigmaAldrich #363073) in a 5:16 v/v ratio, which was sonicated at 65% power for 5 minutes. This second was magnetically stirred at RT for 3 h to evaporate off DCM. Large aggregates were pelleted by centrifugation at 2,000 x *g* for 10 min and supernatants containing the NPs were ultracentrifuged at 80,000 x *g* for 20 min. The NPs were resuspended in DI water and pelleted by ultracentrifugation again. Finally, NPs were resuspended in nuclease-free water, frozen in liquid nitrogen, and lyophilized for 48 h. PLGA NPs were coated with branched PEI (Polysciences, #06090) by reaction with EDC (Thermo Scientific, #22980) and sulfo-NHS (Thermo Scientific, #PG82071) as described in previous works.^26,29^ PLGA-PEI NPs were then washed twice with 1 M NaCl and once with DI water via ultracentrifugation. NPs were resuspended in nuclease-free water and were again lyophilized for 48 h.

The RIG-I agonist PUUC was synthesized according to previously described methods using custom PAGE-purified Ultramer oligonucleotide templates (Integrated DNA Technologies) with the MEGAshortscript™ T7 Transcription Kit (Invitrogen, #AM1354).^27^ PUUC was mixed with PLGA-PEI NPs in 10 mM sodium phosphate buffer (prior treated with diethyl pyrocarbonate (DEPC), pH 6.5) in hydrophobic, low-adhesion tubes under continuous rotation at 4°C for 24 h. After rotation, PUUC loading was confirmed by centrifuging NPs at 10,000 x *g* for 20 min and measuring RNA absorbance in supernatants with the Nucleic Acid Quantification program (Gen 5) using a Synergy HT plate reader and BioTek Take3 microvolume plate. Particle size and surface potential were evaluated with a Zetasizer Nano ZS (Malvern). We have previously reported a PUUC loading efficiency of 100% (5 μg/mg NP) with particle sizes ∼250 nm in diameter with an average surface zeta potential of 31.7 mV and 25.0 mV for PUUC and MPLA+PUUC particles, respectively.^27^

### Intranasal PLP Administration

Female 9-10 wk old Balb/cJ mice were anesthetized with 5% isoflurane. Each treatment group contained three mice. For each group, 4 mg of PUUC NPs (5 μg PUUC per mg NP) or MPLA+PUUC NPs (6 μg MPLA + 5 μg PUUC per mg NP) were resuspended in 60 μL of normal saline (0.9% sodium chloride, Hanna Pharmaceutical, #0409488810) and administered dropwise into the bilateral nares. We have previously confirmed that PLPs are present in the trachea and lungs following intranasal administration using this method.^27^ Mice were euthanized at 4 and 24 h after NP administration to collect lungs. For the RT-PCR experiment, control mice were administered only saline, and lungs were collected at 4 h and 24 h. For the CITE-Seq experiment, naïve mice were used as control.

### RT-PCR Analysis of Lung Gene Expression

After euthanization of mice (n=3 per treatment group) at 4 and 24 h, lungs were collected in 1 mL TRIzol™ Reagent (Invitrogen) using tubes with 1.4 mm ceramic beads. Lung tissues were homogenized in 3 1-minuite increments with a MP Bio FastPrep-24 homogenizer. 200 μL of chloroform was added to the homogenate, and samples were centrifuged for at 12,000 x *g* and 4°C for 15 min. Following mixture separation into a lower red phenol-chloroform phase, interphase, and a clear upper aqueous phase, the aqueous phase was mixed with ethanol in a 1:1 ratio and was added to RNeasy spin columns (Qiagen) for RNA purification. DNAse was added to remove residual DNA, and pure RNA was eluted with RNAse/DNAse free water. RNA yields were quantified with Nanodrop spectrophotometer (Thermo Fisher) and were frozen at 80°C. RNA was used to synthesize cDNA with the SuperScript® III First-Strand Synthesis System (Thermo Fisher) and purified cDNA was kept at -80°C. For each group, cDNA was pooled and 8 ng was combined with SYBR Green Mastermix (Qiagen) for aliquoting into a 384-well RT^2^ Profiler PCR Array with genes from the Mouse Inflammatory Response and Autoimmunity set (Qiagen). Plates were analyzed with the Applied Biosystems™ QuantStudio™ 6 Flex Real-Time PCR System. Within groups, gene expression for each group as normalized to *Actb, B2m, and Gapdh*. Fold regulation scores for each gene per treatment group are provided in **Supplementary Table 1**. Then, each gene per group was normalized relative to the saline control. Upregulated genes with ≥ 1.5 fold change were inputted into the Database for Annotation, Visualization, and Integrated Discovery (DAVID) using the entire 375 microarray gene set as background with DAVID-defined defaults to determine enrichment in categories including those from Gene Ontology (GOTERM), Uniprot (UP_KEYWORDS), INTERPRO, SMART, and KEGG databases. Differential gene expression was visualized on KEGG pathways with the Pathview package.^30^

### CITE-Seq on Lung Cells with the 10x Genomics Platform

We performed CITE-Seq using BioLegend TotalSeq™-A antibodies to generate antibody-derived tags (ADTs) and the 10x Genomics Chromium v3.1 kit to generate single-cell transcripts. Briefly, for three mice in each group and time point, PUUC and MPLA+PUUC PLPs were administered as described above. Naïve mice were used as an untreated control. Mice were euthanized at 4 and 24 h, and lungs were processed into single-cell suspensions with a gentle MACS™ Octo Dissociator and Mouse Lung Dissociation Kit (Miltenyi Biotec, #130-095-297) according to manufacturer protocols.

Cell suspensions were counted and evaluated for viability over 80%. Live cell enrichment (*e*.*g*., Miltenyi Biotech) was processed if the cell viability was less than 80%. Suspensions were centrifuged at 400 x *g* for 5 min and were washed immediately after diluting the samples in 10ml PBS + 0.1% BSA. Cells were counted using Nucleocounter (Cemometec) automated counters. Cells were blocked with 0.5 μL of TruStain FcX™ PLUS (anti-mouse CD16/32) antibody and were incubated at 4°C for 10 min. The antibody cocktail was added to stain the cells and incubated for 30 minutes before proceeding to the wash steps. Three wash steps were performed, followed by filtration. Following cell viability confirmation, the 10x Genomics single cell 3’ v3.1 assay protocol was performed according to the manufacturer’s instructions. To obtain sufficient read coverage for both libraries, we sequenced ADT libraries in 10% and cDNA library fraction at 90% of a lane (Novaseq6000 SP kit).

### CITE-Seq Data Preprocessing

Cells were demultiplexed and aligned using CellRanger v3. ADT reference was used for TotalSeq-A from Biolegend, and reads were aligned to mouse reference genome version mm10-2020-A. Based on transcripts alone, doublet scores were calculated for each cell in each sample individually using the Python-based Scrublet package.^31^ ADTs were normalized and denoised with the dsb method according to the originally published method using CellRanger-defined cell identities followed by filtering for outlying cells with more than ± 3 times the mean absolute deviation of total RNA and ADT counts and with a mitochondrial DNA percentage of less than 25%.^32^ Background for dsb normalization was defined as droplets with ADT counts between 1,500 and 2,500 and less than 100 genes recorded; isotype controls were used from the TotalSeq-A set (IgG1, IgG2, IgG2b). Cells were preliminarily clustered using ADTs only to assess normalization quality.

Cells were further preprocessed using Seurat V4 to filter out cells containing less than 200 total RNA counts and more than 10% globin genes. For each sample, gene expression matrices were normalized based on 4000 features with the SCTransform method while regressing mitochondrial counts.^33^ Cell cycles scores for S and G2M phases were calculated with mouse homologs of the gene list from Tirosh et al. (2019 update).^34^ For cell identification, SCT-normalized transcript counts were integrated with Seurat for the top 6000 variable genes using the top 3000 variable genes as anchors, and effects due to cell cycle scores were regressed out.^35^ ADTs were integrated based on dsb-normalized values neglecting isotype control reads. Cells were clustered based on both RNA and ADT integrated matrices via the weighted nearest neighbors (WNN) approach. Marker genes were preliminarily assigned, and metaclusters were subclustered (i.e. T/NK cells, macrophage/monocytes/DCs, etc. were subsetted and analogously reclustered) to find small doublet clusters. Small clusters expressing arrangements of marker genes from multiple lineages (i.e. *Cd3e* and *Cd19* and *Vwf*) or high propensity of mitochondrial genes as markers were filtered out of the data. Additionally, ADTs were manually visualized for poor staining, with 68 ADTs excluded from further analysis. All ADTs, including those excluded with reasons given, are provided in **Supplementary Table 2**. ADTs and transcripts were matched based on the analysis of Pombo et. al.^36^ Our analysis yielded 25,439 remaining cells and 18,010 unique genes across all treatment and control groups (**Table 1**). These samples were analogously integrated, clustered, and subclustered in the absence of confounding doublets and low-quality cells to yield the final cohort of cells for further biological analysis.

**Table 1.** Sequencing Summaries of Sample Reads and Processed Cell Counts. Mean reads and median genes are given on a per-cell basis as reported from the CellRanger output. The estimated Cell Count represents the contents of filtered matrices prior to further processing in Seurat. Processed Cell Count represents the cells used for further biological analysis.

| Sample | Mean Reads / Cell | Median Genes / Cell | 10X Estimated Cell Count | Processed Cell Count |
| --- | --- | --- | --- | --- |
| naive | 37701 | 1692 | 3109 | 2588 |
| p4 | 20407 | 1254 | 7784 | 6586 |
| mp4 | 21104 | 1234 | 8838 | 7753 |
| p24 | 32769 | 1268 | 4752 | 3994 |
| mp24 | 32376 | 1120 | 5086 | 4518 |
| **Totals:** |  |  | **29569** | **25439** |

### Differential Gene Expression (DEG), Gene Expression Scoring, and Gene Set Enrichment Analysis (GSEA)

Cluster marker genes were calculated with Seurat using a Wilcoxon Rank Sum (WRS) test on SCTransformed transcripts for each integrated WNN-defined cluster against all other clusters. Cell types were assigned according to known scRNA-seq cell atlases.^10,37,38^ A final list of cell type markers across all treatments and controls is provided as **Supplementary Table 3**. Differential gene expression (DEGs) within cell types between treatment groups was generated with a WRS test. Supplementary files are provided for DEGs from all treatment groups compared to naïve (**Supplementary Table 4**) and for dual treatment groups (MPLA + PUUC) compared to single treatment (PUUC) for each cell type (**Supplementary Table 5**). For subclustered neutrophils, gene expression scores were calculated with AddModuleScore in Seurat based on mouse gene lists for ‘Positive regulation of apoptotic process (GO:0043065),’ ‘Necroptosis (GO:0070266),’ ‘Chemotaxis (GO:0030593),’ ‘Phagocytosis (GO:0006911),’ and ‘NADPH oxidase (Henderson and Chappel, 1996)’ as reported by Xie et al.^39,40^ IFN-γ responsive scores were calculated likewise with the Hallmark Interferon Gamma Response gene list.^41^ Gene set enrichment analysis (GSEA) was performed on DEGs expressed in at least 5% of cells for a given cell type-treatment-timepoint combination using the fgsea package in R with a minimum bin size of 15 and no maximum.^42^ Mouse gene lists from Hallmark sets, KEGG pathways, Gene Ontology Biological Process (GOBP), and Reactome were used for GSEA as provided by the msigdbr package using the Molecular Signature Database v7.4.^41^ The results of significant GSEA enrichments for the treatment groups vs. naïve and MPLA + PUUC vs. PUUC comparisons are provided as **Supplementary Table 6** and **Supplementary Table 7** respectively.

### RNA Velocity Analysis

Spliced and unspliced transcript matrices were generated from the 10X Chromium filtered output with velocyto using Python.^43^ Based on the previous clustering, transcripts corresponding to neutrophils were subsetted. Neutrophil transcripts were processed into RNA velocity estimates with scVelo using dynamical modeling.^44^ For matrices corresponding to 4 hr treatments, a root cell was manually defined based on high expression of early neutrophil developmental genes (i.e. *Camp*) as visualized with UMAP of unintegrated transcripts in Seurat. Based on these dynamics, latent time for each neutrophil within each treatment group was calculated and visualized on the Seurat-generated UMAP coordinates. Genes expressing the highest dynamical behavior across latent time were ranked and visualized with phase portraits.

### Intercellular Communication Analysis

Based on Seurat-defined clusters, cell types with less than 10 cells in at least one treatment were removed for communication analysis. For transcript matrices for each treatment group, predefined pathways including “Secreted Signaling,” “ECM Receptor,” and “Cell-Cell Contact” were used to compute probable cell-cell interaction networks with the CellChat package in R 4.0.1.^45^ Default settings for unsorted single cells were used. Statistically significant interactions were visualized with circle and chord plots.

## RESULTS

### PUUC and MPLA+PUUC nanoparticles alter cell-type proportions

Nanoparticles were synthesized to deliver PUUC and MPLA to mouse lungs. Briefly, PLGA nanoparticles were synthesized by water-oil-water antisolvent evaporation with and without encapsulated MPLA. Surfaces were made positively charged via amidification with bPEI, enabling electrostatic loading of PUUC mRNA (**Fig. 1A**).

**Figure 1.**
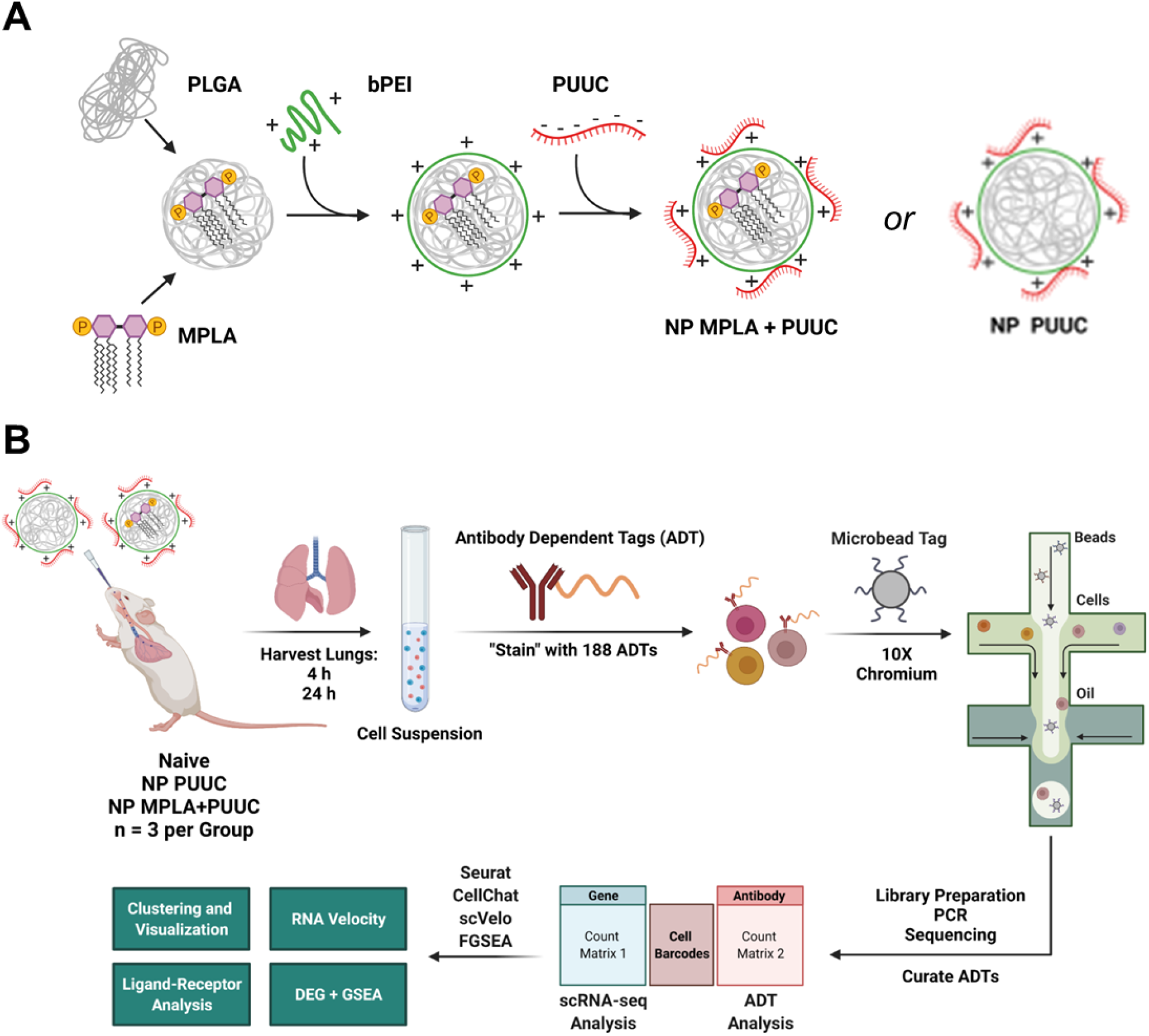
Synthetic, experimental, and analytical overview for PUUC and MPLA+PUUC nanoparticle stimulation at early time points in the murine lung. **A)** Schematic of nanoparticle (NP) synthesis. PLGA NPs with or without mono-phosphoryl lipid A (MPLA) were first synthesized by double-emulsion antisolvent evaporation, followed by covalent modification with branched polyethyleneimine (PEI) with EDC/sulfo-NHS to make positively-charged PLGA-PEI NPs. Negatively charged PUUC RNA was then electrostatically loaded onto the particles to make either PUUC or MPLA+PUUC NPs. **B)** Schematic of experimental and analytical overview. 4 mg of PUUC and MPLA+PUUC NPs were combined with 60 μL saline and were administered into the bilateral nares of three mice per group. Doses of 20 ug PUUC and 24 ug MPLA were used per mouse. Lungs from these groups and naïve, untreated mice were harvested, processed into single-cell suspensions, and were pooled groupwise at 4 and 24 hours. Suspensions were mixed with DNA-barcoded antibodies (antibody-dependent tags; ADTs) and were processed according to 10X Chromium Protocols. cDNA libraries were prepared by reverse transcription, amplified with PCR, and sequenced. After alignment, matrices of gene (RNA) and protein (ADT) expression counts were processed in R 4.1.0 with Seurat, CellChat, scVelo, and FGSEA for clustering and visualization, ligand-receptor analysis, RNA velocity computation, and differential gene (DEG) expression followed by Gene Set Enrichment Analysis (GSEA).

PUUC or MPLA+PUUC NPs were intranasally administered to mice, and lungs were harvested at 4 and 24 hours to evaluate immunostimulatory potential, RT-PCR microarray analysis yielded a broad set of DEGs in each group (**Fig. S1A**). Functional enrichment analysis showed that upregulated DEGs, especially at 24 hours, were preferentially associated with secreted products like cytokines and chemokines, particularly those carrying the C-X-C motif (**Fig. S1B**). Interestingly, PUUC, a RIG-I agonist, alone was associated with the upregulation of the KEGG TLR signaling pathway at 24 hours (**Fig. S1C**). This overlap between RIG-I and TLR signaling motivated the further study of PUUC, combined with the TLR4 agonist MPLA in murine lungs.

After intranasal administration of PUUC and MPLA+PUUC NPs, lungs were harvested at 4 and 24 hours, and tissues were processed for CITE-Seq (**Fig. 1B**). After preprocessing, cells from each group were integrated and clustered based on combined gene and surface marker expression (**Fig. 2A**). Independent of gene expression, surface CD45 readily separated immune cells from structural cells expressing ESAM or EPCAM (CD326). CD11b identified monocytes and granulocytes, Ly-6G for granulocytes, CD19 for B cells, and SiglecF (CD170) for alveolar macrophages (AMs). T cells could be identified based on TCRB expression followed by differentiating between CD4 and CD8 (**Fig. 2C**). Gene expression helped narrow further the expression of other cells including classical monocytes (*Ly6c2*), nonclassical monocytes (*Gngt2*), interstitial macrophages (*Apoe, C1q* genes), pDCs (*Siglech*), neutrophils (*S100a8/9, Retnlg*), gCap endothelial cells (*Gpihbp1*), aCap endothelial cells (*Kdr*), vein endothelial cells (*Vwf*), AT1 cells (*Hopx*), AT2 cells (*Sftpc*), myofibroblasts (*Tagln*), *Ebf1*+ fibroblasts (also expressing *Postn*), lipofibroblasts (*Inmt*), basophils (*Cyp11a1, Mcpt8*), adventitial fibroblasts (*Dcn*), lymphatics (*Ccl21a*), ciliated cells (*Scgb1a1*), NK cells (*Gzma*), and nuocytes/ILC2s (*Rora*), regulatory CD4 T cells (*Ctla4*), gamma-delta T cells (*Trdc*), and a population of proliferating T cells (*Ctla4, Pclaf, Top2a*) (**Fig. 2D, Supplementary Table 3**). B cells were identified based on surface expression of CD19 and expression of immunoglobulin transcripts like *Igkc* with mature B cells expressing *Ighm* and surface IgD and immature B cells expressing *Ighm* in the absence of IgD (**Fig. S2A-C**).^46^ Cell type identities were largely assigned according to known scRNA-seq-based markers, except for two small populations of dendritic cells. CD103 DCs highly expressed surface CD103 and *Ccl17* and *Ccl22* and H2 Hi DCs expressed MHC-II transcripts (*H2-DMb1, H2-Eb1, H2-Aa, H2-Ab1, Cd74*) in the absence of surface CD103 (**Supplementary Table 3**). At 24 hours, neutrophils were found to be the most populous cell type in the PUUC- and especially the MPLA+PUUC-stimulated lungs (**Fig. 2B**).

**Figure 2.**
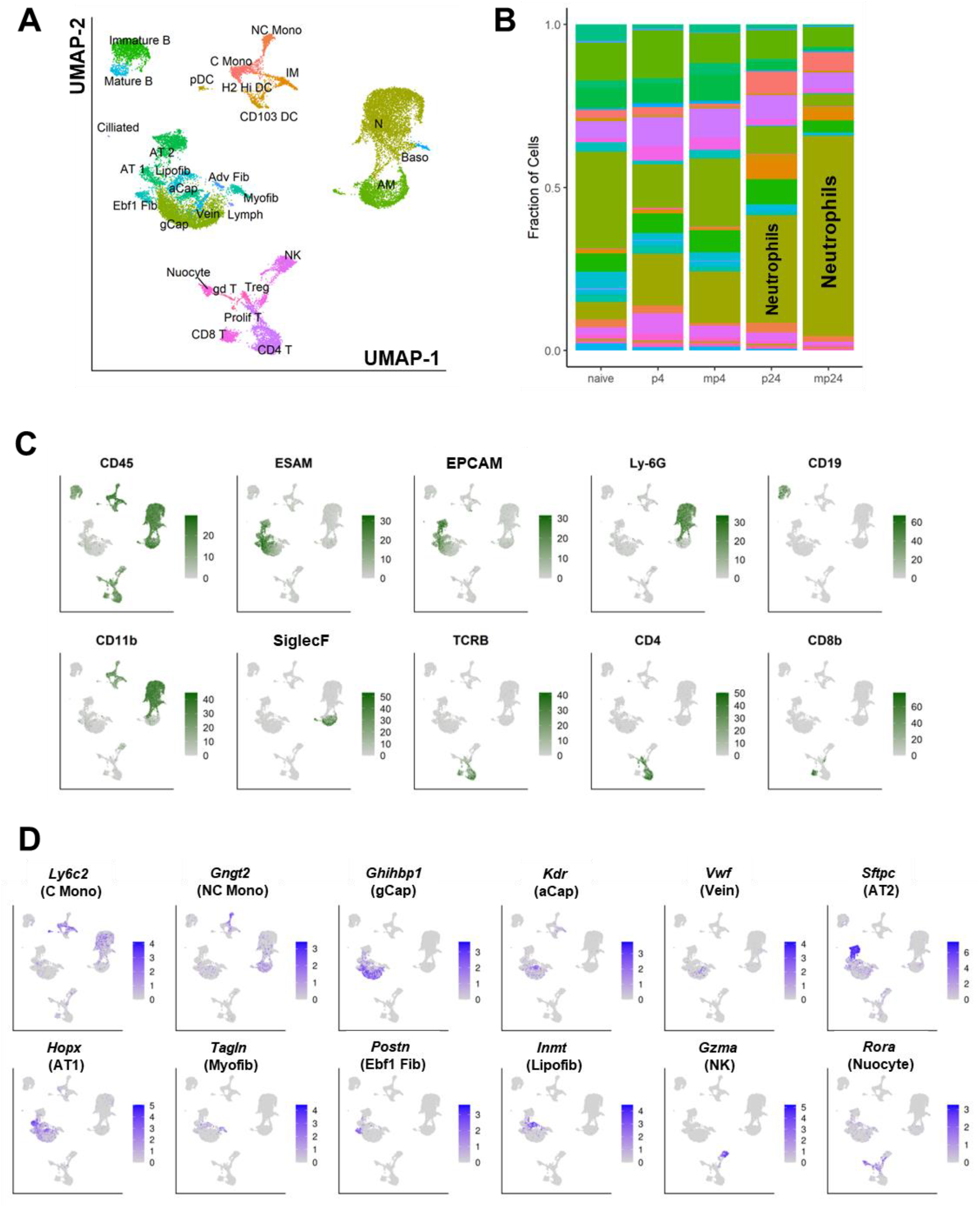
CITE-Seq identifies differentially populous cell types responding to PUUC and MPLA+PUUC stimulation. **A)** Gene and protein expression matrices from the naïve, P4, MP4, P24, and MP24 groups were integrated and clustered together using weighted-nearest-neighbor (WNN) uniform manifold approximation projection (UMAP). Graph-based clusters were identified based on expression of known marker genes and surface protein expression. **B)** After cluster identification, cell types as a fraction of total cells in each group were visualized. **C)** Surface expression of 10 predominant ADTs were expressed on the original UMAP projection, as were the **D)** genes expressed for 12 predominant cell types. ADT expression values were normalized with DSB. UMAP and corresponding plots of surface protein and gene expression were generated with Seurat V4. All figures were generated using R 4.1.0.

### Interferon-stimulated neutrophils differentially regulate vesicular transport and cytokine/chemokine expression in PUUC- and MPLA+PUUC-stimulated lungs

Given the increase in neutrophil cell population in the 24-hour treatments, we subclustered neutrophils for further analysis without correcting for treatment effects. On UMAP, neutrophils from the 4 h, 24 h, and naïve groups clustered in three distinct groups (**Fig. 3A**). Independent clustering of neutrophils revealed 4 distinct clusters with N0 and N2 comprised mainly of 24-hour neutrophils (from either the PUUC or MPLA+PUUC treatments), N1 comprised of 4-hour neutrophils, and N3 comprised of neutrophils from the naïve lung (**Fig. 3B**).

**Figure 3.**
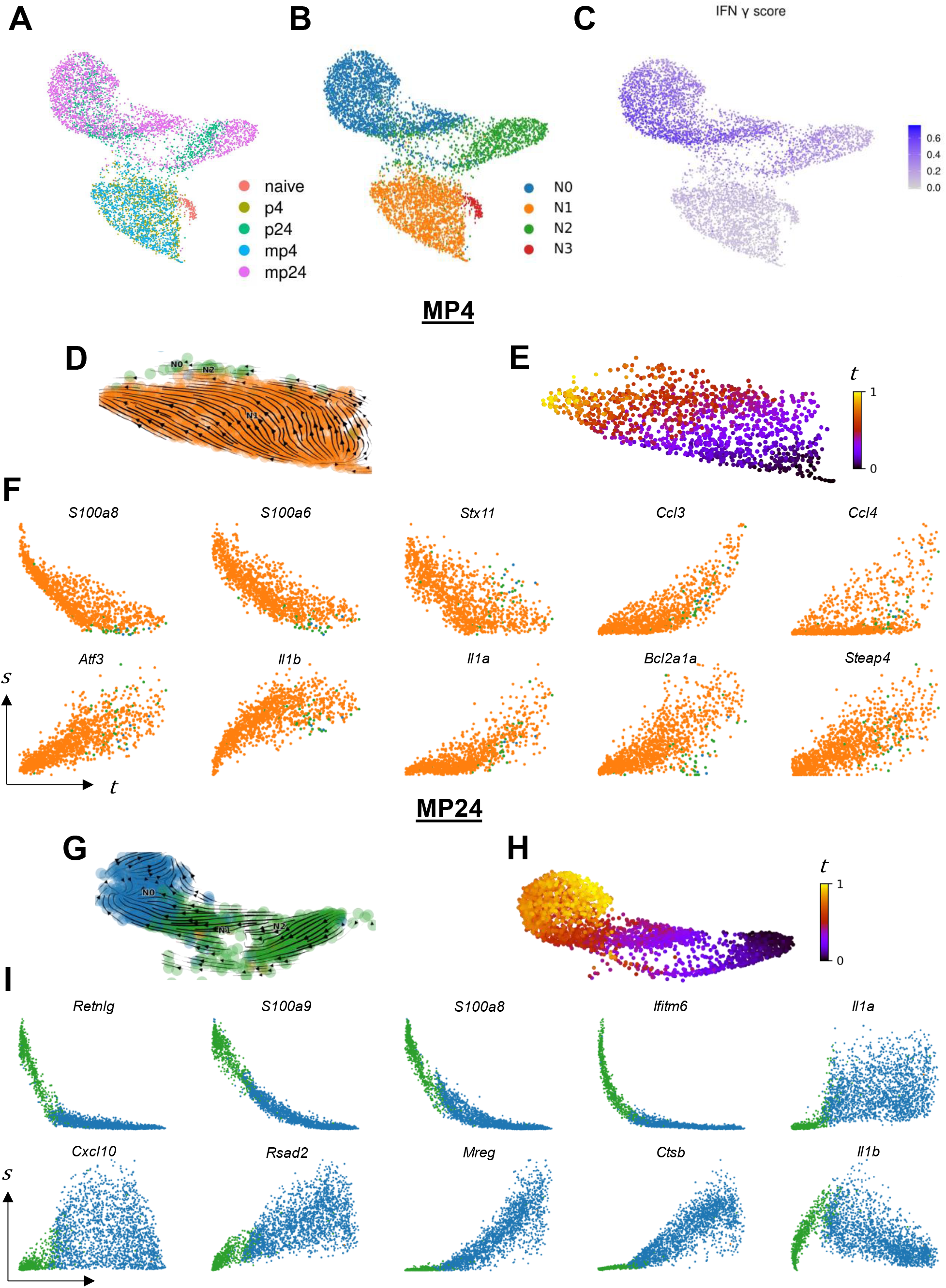
Trajectory analysis of invading neutrophil gene expression along latent time from MPLA+PUUC stimulated lungs at 4 and 24 hours. **A)** Neutrophil gene expression data for all treatments was visualized with UMAP without integration, and cells were labeled by treatment group. **B)** Gene expression was analyzed by Louvain clustering to identify four neutrophil subclusters. **C)** Cell expression of the 200 genes in the MSigDB v7.4.1 HALLMARK_INTERFERON_GAMMA_RESPONSE category were scored with the AddModuleScore function. **D)** Spliced and unspliced transcript counts were computed for all cells in each treatment group independently with velocyto. Neutrophils from the MPLA+PUUC 4hr (MP4) treatment group were subsetted from other cell types, and RNA velocity profiles were computed with scVelo using dynamic modeling. RNA velocity embeddings were visualized on the original UMAP. **E)** Based on RNA velocity modeling, neutrophils were placed into a continuous latent time, *t*. **F)** Genes were selected from a group of the top 15 most-likely driver genes in the dynamic model, and expression dynamics of spliced transcripts, *s*, were visualized across latent time. Panels **G-I**) Indicate the same analysis for neutrophils from the MPLA+PUUC 24h (MP24) treatment group. Colors in phase portraits correspond to cluster colors in panel B. UMAP coordinates, module scores, and clustering were computed with Seurat V4 in R 4.1.0. veclocyto and scVelo were run with Python 3.7.

N3 preferentially expresses genes related to respiratory burst, including NADPH Oxidase Activator 2 *(Ncf2)*, and cell structure and motility, including actin transcripts (*Actb* and *Actg1*) and the actin regulator Thymosin Beta 4 (*Tmsb4x*). Additionally, N3 is enriched for ribosome-related genes like *Rps29, Rps27*, and *Rpl37* (**Fig. S3A**). GSEA on DEGs from the N1 vs. N3 comparison confirm enrichment in categories related to respiratory burst, cytoskeletal organization, and especially the ribosome (**Supplementary Table 8**). N1 primarily expresses the C-X-C and C-C motif cytokine transcripts *Cxcl2, Cxcl3, Ccl3*, and *Ccl4*. GSEA confirms enrichment in the reactome categories “Peptide ligand binding receptors” and “Chemokine receptors bind chemokines” in N1 (**Supplementary Table 8**). Within the 24 h neutrophil groups, N2 expresses Ca-sensing alarmins like *S100a8* and *S100a9*, interferon-induced genes like *Ifitm1* and *Ifitm6*, and neutrophil secondary granule components such as *Ngp* and *Camp*. In contrast, N0 highly transcribes cathepsins like *Ctsb* and *Ctsz*, MHC-I related genes *B2m* and *H2-K1*, and cytokines like *Csf1* (M-CSF) and *Ccl3* (**Fig. S3B**). Cells in N0 score higher in expression of IFN-γ-related genes (**Fig. 3B**), and GSEA on DEGs from these cells are preferentially enriched in the reactome category “interferon gamma signaling.” (**Fig. 3C, Supplementary Table 9**). While the 24 h neutrophils in N0 and N2 score higher in apoptosis and necroptosis gene expression scores, the difference in these scores between the N1 and N3 clusters do not appear as pronounced as those between IFN-γ scores (**Fig. S3F-H**).

Given that some cells within each the 4 and 24 h neutrophil clusters express genes associated with neutrophil maturation in the bone marrow like *Camp, Ngp*, and *Mmp8* (**Fig. S3C-E**), we hypothesized that neutrophils in these data may capture a continuum of neutrophil development in the context of PUUC- and MPLA+PUUC-induced inflammation. We therefore computed the RNA velocity profiles of neutrophils in each treatment group independently (**Fig. 3D, 3G, S4A, S4D**). Cells from each computation were fit to a latent time model **(Fig. 3E, H, S4B, S4E)**. The top 15 genes associated with the latent time trajectory were assessed. At 4 hours, PUUC-induced neutrophils show signs of increasing TNFa-related NF-kB signaling with increasing *Nfkbia* and *Tnfiap3* expression over latent time **(Fig. S4C)**. Neutrophils from the MPLA+PUUC 4-hour also showed induction of the TNFa-induced *Steap4* the induction of the master cell stress regular *Atf3* **(Fig. 3F)**. For both 4-hour groups, neutrophils increased synthesis of CC chemokines and IL-1-related transcripts like *Ccl4, Il1a*, and *Il1b* while also decreasing expression of the vesicular transport protein Syntaxin 11 *(Stx11)*. In the PUUC 24-hour group, neutrophils increased synthesis of M-CSF (*Csf1*) and the IL-1 receptor antagonist (*Il1rn*). In the same group, the mouse-specific genes *Wfdc17* and *Trim30c* were increased and decreased, respectively **(Fig. S4F)**. In the MPLA+PUUC neutrophils at 24 hours, *Ifitm6* decreased over latent time, while analogously the interferon-stimulated antiviral protein Viperin (*Rsad2*) was increased. The vesicular transport protein melanoregulin (*Mreg*) and lysosomal protease cathepsin B (*Ctsb*) were also increased in this group. In both 24-hour groups, *Il1b* increased and then decreased over latent time. Overall, neutrophils from each treatment and both timepoints showed a common decrease in genes from the S100A family of proteins (i.e. *S100a8*).

### PUUC and MPLA+PUUC induce a broad decrease in ribosomal protein expression within 4 hours of stimulation within multiple cell types

Given the changes in neutrophil activity in the treated lungs, we computed DEGs for all cell types and treatment combinations compared to naïve lungs (**Supplementary Table S4**). GSEA on DEGs was computed, and gene sets related to ribosome function was found to be significantly downregulated in nearly all cell types at 4 hours including GOBP_TRANSLATION_INITIATION, KEGG_RIBOSOME, and REACTOME_TRANSLATION (**Fig. S5A**). Only NK cells at from both treatments and aCap endothelial cells from PUUC-treated lungs were found to not be significantly downregulated at 4 hours (**Fig. 4A**). Instead, these aCap endothelial cells were found to be significantly upregulating metallothioneins (*Mt1, Mt2*), cytokines (*Cxcl2, Il1b*), and the A-kinase anchoring protein 12 (*Akap12*) (**Supplementary Table S4**). *Akap12* is particularly associated with bringing components of the Protein Kinase A (PKA) signaling cascade to the cell membrane, suggesting a role for PKA signaling in early endothelial cell-mediated inflammation.^47^

**Figure 4.**
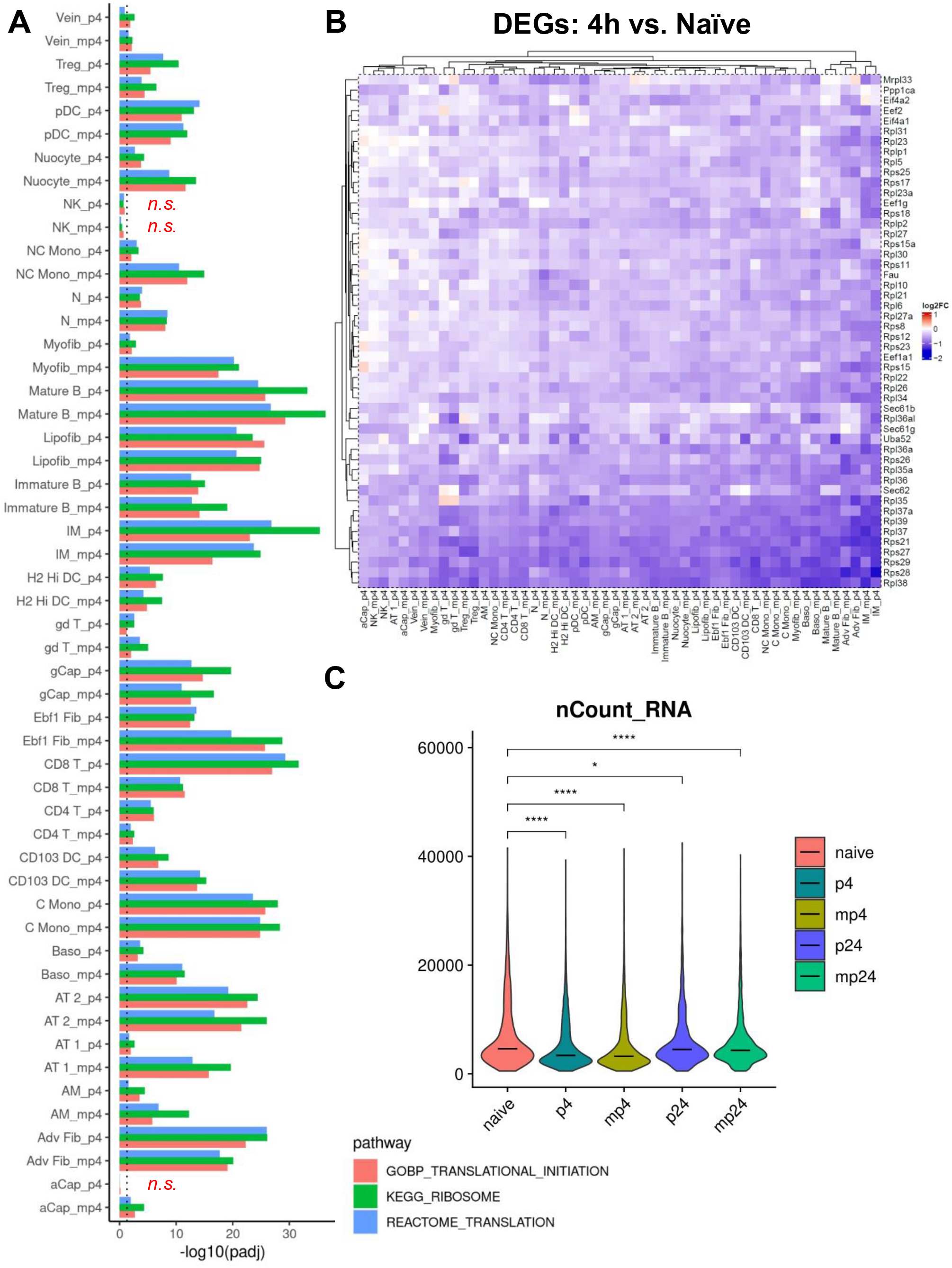
PUUC and MPLA+PUUC induce a near-global decrease in ribosomal protein expression within 4 hours. **A)** Negative enrichment of select ribosome-related gene sets (GOBP_TRANSLATION_INITIATOIN, KEGG_RIBOSOME, and REACTOME_TRANSLATION) was compared between lung cell types by significance. Dotted line represents the cutoff of significant enrichment (P_adj_ = 0.05). **B)** Heatmap of the 50 most downregulated genes in the KEGG, REACTOME, and Gene Ontology categories from Figure S5 with independent hierarchical clustering of genes and cell types in the 4-hour treatment groups. **C)** Total RNA counts for all genes in each cell was visualized with violin plots for each treatment group with a horizontal line representing median expression. For comparisons between transcript counts, *P < 0.05, ****P < 0.0001 as determined by Kruskal-Wallis test. Heatmap generated with ComplexHeatmap package in R. All figures were generated with R 4.1.0.

The top 50 most downregulated ribosome-related genes included components of the small (*Rps28*) and large ribosomal subunit (*Rpl38*), elongation factors (*Eef1a1*), and components of the ER translocation apparatus (*Sec62*) (**Fig. 4B**). In contrast, cells at 24 hours were less uniformly associated with downregulation in ribosomal gene sets (**Fig. S5B**). In addition to decreases in translational genes, overall transcript counts were decreased at 4 hours in cells from PUUC- and MPLA+PUUC-stimulated lungs indicating an increase in mRNA turnover and/or a decrease in global transcription (**Fig. 4C**).

In the absence of significant ribosomal gene downregulation, NK cells were associated with positive differential gene expression associated with NF-kB activation including *Nfkbia* and *Nfkbiz*, cell proliferation and survival (*Pim1*), and cell stress (*Gadd45b*) in NK cells from both PUUC and MPLA+PUUC lungs at 4 hours (**Fig. S6A-B**). Indeed, GSEA on DEGs from both 4-hour NK cell groups were similar, showing similar positive enrichment in categories associated with the P38-MAPK cascade, transcription factor activity, lipid storage, and response to biotic stimuli (**Fig. S6C**). In both groups, the top positively enriched categories were associated with leading edges that included *Gadd45b, Nfkbia, and Nfkbiz* whereas downregulated categories were associated with cell-surface function with leading edges predominately featuring integrin subunit beta 2 (*Igtb2*) (**Fig. S6D**).

### PUUC and MPLA+PUUC are associated with widespread interferon-related transcriptional responses in the lung at 24 hours

In contrast to the widespread decrease in ribosome-associated genes at 4 hours, we found a widespread increase in interferon-responsive genes at 24 hours in cell types from both the PUUC- and MPLA+PUUC-treated lungs. For all cell types, GSEA on DEGs showed an enrichment in IFN-related categories including, HALLMARK_INTERFERON_ALPHA_RESPONSE and HALLMARK_INTERFERON_GAMMA_RESPONSE (**Fig. 5A**). The same broad increase was not seen at 4 hours, but we do note that adventitial fibroblasts and interstitial macrophages were particularly enriched in interferon-related categories compared to other cell types at 4 h (**Fig. S7A-B**). These gene sets were predominated by C-X-C and C-C cytokines and metallothionein 2 (*Mt2*) **(Supplementary Table S6)**. Likewise, at 24 h, genes from these categories included C-X-C and C-C chemokines like *Cxcl10* and *Ccl4*, interferon stimulated genes (*Isg15*) and their inhibitors (*Usp18*), intracellular antiviral proteins including Viperin (*Rsad2*) and Z-DNA binding protein 1 (*Zbp1*), transcription factors like *Irf7*, and the MHC-I component β_2_ microglobulin (*B2m*) (**Fig. 5B**). According to a transcript-based intercellular communication model generated with CellChat, IFN-β was predicted to be secreted by NK cells, nuocytes, interstitial macrophages, DCs, Tregs, and CD8 and CD4 T cells in both PUUC and MPLA+PUUC-stimulated lungs at 24 hours, with aCap endothelial cells being an additional *Ifnb1* synthesizer in the MPLA+PUUC group (**Fig. 5C**). Type II interferon transcripts were less extensively detected in the dataset, with only NK cells showing significant IFN-γ (*Ifng*) signaling to CD4 T cells, H2 Hi DCs, classical monocytes, and basophils in the PUUC 24 h group and to CD4T cells, nonclassical monocytes, adventitial fibroblasts, and aCap endothelial cells in the MPLA+PUUC 24 hour group (**Fig. 5D**). It should be noted that these cell-cell interactions only consider the transcripts of ligands and receptors and may underestimate signaling between ligands and cell-surface receptors not undergoing active transcription.

**Figure 5.**
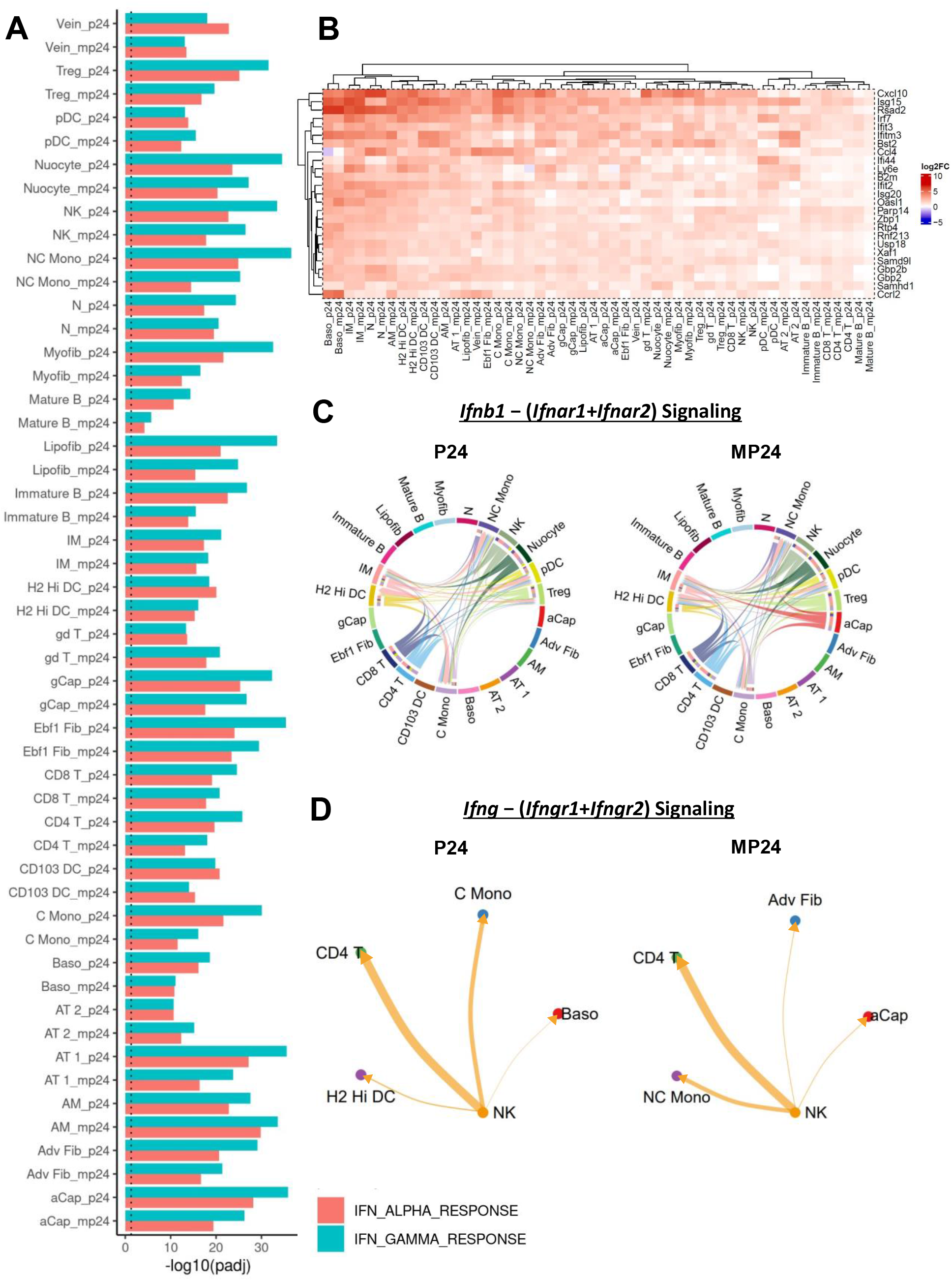
Diverse cell types express interferon-responsive genes in the PUUC and MPLA+PUUC treated lungs at 24 hours. Gene set enrichment analysis (GSEA) for differentially expressed genes (DEGs) were computed with FGSEA for all cell types and treatment groups compared to the naïve group. **A)** Positive enrichment for the HALLMARK_IFN_ALPHA_RESPONSE and HALLMARK_IFN_GAMMA_RESPONSE categories was visualized with a bar plot for all cell types at 24 hours. Predicted interactions between **B)** *Ifnb1* − (*Ifnar1*+*Ifnar2*) and **C)** *Ifng* − (*Ifngr1*+*Ifngr2*) were visualized with chord diagrams and circle plots, respectively. **D)** A heatmap was used to visualize expression of the top 25 DEGs vs. naïve from an aggregate of genes in the following gene sets: HALLMARK_INTERFERON_GAMMA_RESPONSE, GOBP_RESPONSE_TO_INTERFERON_GAMMA, HALLMARK_INTERFERON_ALPHA_RESPONSE, REACTOME_INTERFERON_SIGNALING, and REACTOME_INTERFERON_ALPHA_BETA_SIGNALING. Dotted line represents a significance of P = 0.05. Punitive cell-interaction networks based on transcriptome profiling was computed with CellChat for all experimental groups. Only significant *(P < 0*.*05)* predicted cell interactions were shown. Heatmap generated with the ComplexHeatmap package. All figures were generated with R 4.1.0.

### The addition of MPLA to PUUC preferentially favors neutrophil chemotaxis, ion homeostasis, and TNFa-related signaling in multiple cell types

To determine differences between the treatment groups, DEGs were computed between the MPLA+PUUC vs. PUUC-simulated groups at 4 and 24 hours (**Supplemental Table S5**). Compared to the PUUC-stimulated group, GSEA on DEGs showed positive enrichment in many cell types in many proinflammatory categories including REACTOME_IL10_SIGNALING, HALLMARK_TNFA_SIGNALING_VIA_NFKB, GOBP_NEUTROPHIL MIGRATION, and GOBP_TRANSITION_METAL_ION_HOMEOSTASIS (**Fig. S8**). Alveolar macrophages were among the most enriched cell types in these categories at both 4 and 24 hours in the MPLA+PUUC group. DEGs in AMs from 4 hours included transcripts for cytokines like *Tnf, Il1a*, and *Il1b*, C-C and C-X-C chemokines like *Ccl4* and *Cxcl1*, and hepcidin (*Hamp*). Signs of NF-kB and TNF-a stimulation were indicated by expression of *Nfkbia* and *Tnfaip2*, respectively (**Fig. 6A**). At 24 hours, AMs were upregulated in DEGs related to ion-binding function including the antimicrobial iron-sequestering protein lipocalin 2 (*Lcn2*), ferritin heavy subunit 1 (*Fth1*), metallothionein 2 (*Mt2*), and calcium/manganese-binding S100 proteins (*S100a8, S100a9*) (**Fig. 6B**). Generally, C-C and C-X-C chemokines (especially a trio of cytokines: *Cxcl2, Ccl3, Ccl4*) and metal-binding proteins (such as haptoglobin, *Hp*) were upregulated in multiple other cell types, including *Ebf1*+ fibroblasts, vein endothelial cells, adventitial fibroblasts, and AT1 cells at 24 hours (**Fig. 6C**). Given the increase in general C-X-C chemokines in the MPLA+PUUC treatment group, we visualized predicted CXCL1 signaling in the PUUC and MPLA+PUUC lungs at 24 hours. In PUUC lungs, AMs were the most predominant cell type transcribing *Cxcl1* which was predicted to stimulate neutrophils expressing the receptor, *Cxcr2*. Structural cells such as adventitial fibroblasts, myofibroblasts, and AT2 endothelial cells also contributed to CXCL1 signaling to neutrophils in PUUC lungs (**Fig. 6D**). In the dual-treatment group, MPLA+PUUC was associated with the same *Cxcl1-Cxcr2* interactions with the addition of Ebf1+ fibroblasts, CD103+ DCs, and aCap endothelial cells contributing to *Cxcl1* transcription (**Fig. 6E**). As CXCL1 is a neutrophil chemoattractant, this increase in *Cxcl1* production may contribute to the overall increase in neutrophils in the MPLA+PUUC lungs at 24 hours.

**Figure 6.**
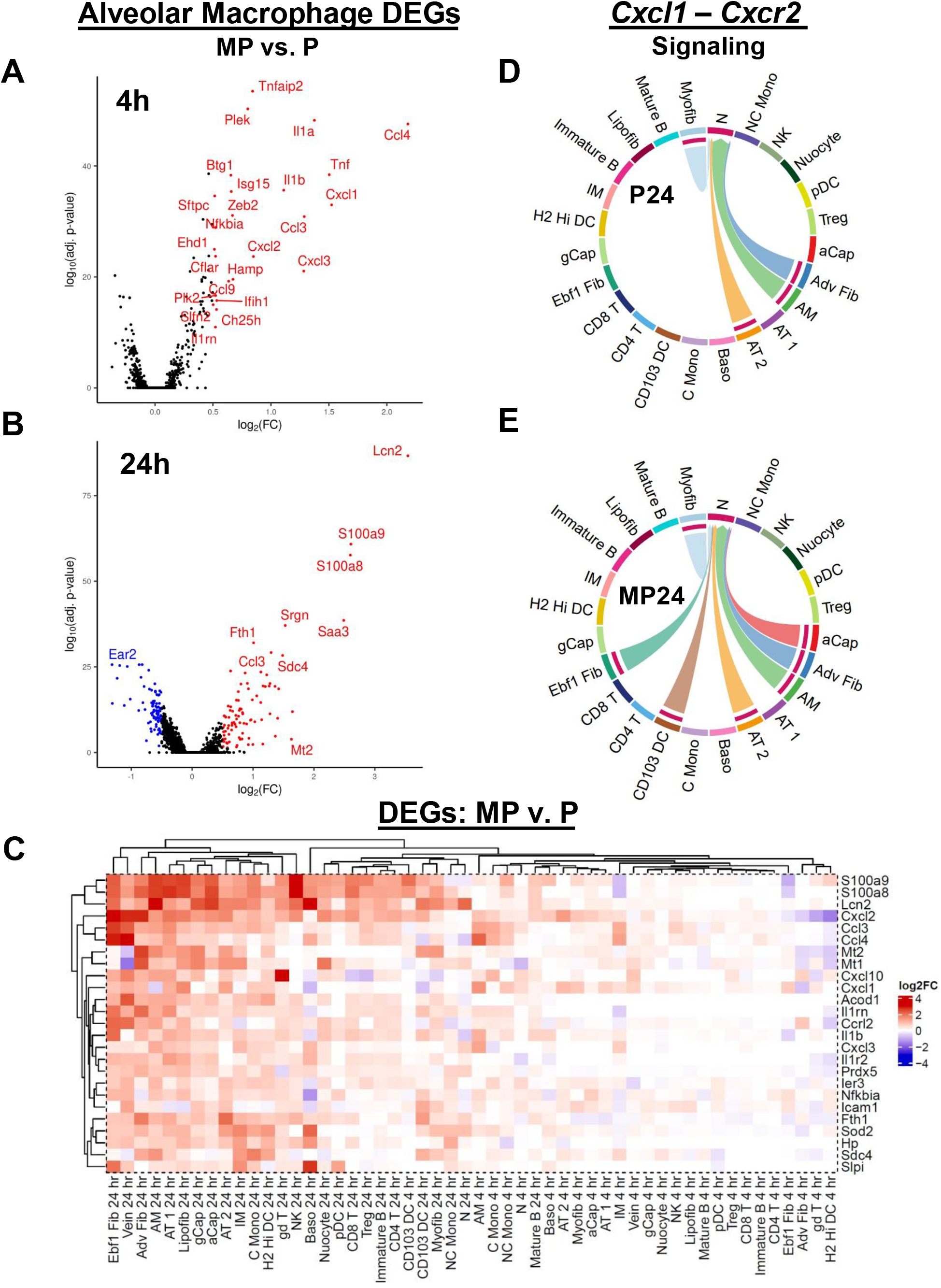
MPLA+PUUC lung stimulation differentially favors neutrophil chemotaxis, ion homeostasis, and TNFa-related signaling compared to PUUC stimulation alone. Differentially expressed genes (DEGs) from alveolar macrophages between the MPLA+PUUC vs. PUUC treatment were visualized for the **A)** 4 and **B)** 24 h timepoints with volcano plots. **C)** The top 25 DEGs from gene sets related to neutrophil chemotaxis, transition metal ion homeostasis, and TNFa-related signaling were depicted on a heatmap. Predicted signaling interactions between *Cxcl1* and *Cxcr2* were computed **D)** for PUUC and **E)** MPLA+PUUC at 24 h across all cell types and were visualized with Chord diagrams. Genes from the following GSEA categories were aggregated to produce the heatmap: GOBP_NEUTROPHIL_MIGRATION, GOBP_GRANULOCYTE_CHEMOTAXIS, GOBP_NEUTROPHIL_CHEMOTAXIS, GOBP_GRANULOCYTE_MIGRATION, GOBP_LEUKOCYTE_CHEMOTAXIS, GOBP_CELLULAR_TRANSITION_METAL_ION_HOMEOSTASIS, GOBP_TRANSITION_METAL_ION_HOMEOSTASIS, HALLMARK_TNFA_SIGNALING_VIA_NFKB, REACTOME_INTERLEUKIN_10_SIGNALING, GOBP_INFLAMMATORY_RESPONSE, GOBP_RESPONSE_TO_MOLECULE_OF_BACTERIAL_ORIGIN. DEGs were computed with Seurat. Heatmap was produced with ComplexHeatmap. Predicted cell interactions were generated with CellChat, and only significant interactions are shown. All analysis was generated with R 4.1.0.

## DISCUSSION

We have previously reported that RIG-I and TLR4+RIG-I with antigen can induce adaptive immune responses in the mouse muscle and lung.^27^ Here, we characterize the early innate immune response to the RIG-I agonist PUUC and dual RIG-I+TLR4 agonist combination MPLA+PUUC in the lung alone to understand early events associated with RIG-I and TLR4 signaling which are common in lung pathogens, as well as identify whether such nanoparticle-PAMPs could be used for immunotherapies or vaccines in the future. We employed nanoparticles to deliver these PAMPs to the lung since RLRs are cytoplasmic receptors and the RIG-I agonists are large, charged molecules unable to enter cells without a carrier. Furthermore, co-delivery in particles increases the chance of stimulating the same cell with both ligands, similarly to a pathogen, which are also particulate carriers with multiple PAMPs.

Our data demonstrate that multiple processes occur in structural and hematopoietic cells, including the downregulation of ribosomal protein genes at 4 h and upregulation of antiviral genes at 24 h. These results may relate to the previous findings that pediatric influenza patients rely on combined TLR4 and RIG-I functions to prevent organ dysfunction and respiratory failure following influenza-related critical illness.^48^ Ribosome-associated transcripts are especially decreased in adventitial fibroblasts, interstitial macrophages, AT2 epithelial cells, classical monocytes, and B cells in both treatment groups. Adventitial fibroblasts and interstitial macrophages are also cells that display early synthesis of ISGs at 4 h that foreshadows the ubiquitous expression of ISGs in all cells at 24 h. The most highly expressed components of these gene sets include C-C and C-X-C cytokines that promote neutrophil chemotaxis and metallothionein that increase intracellular zinc sequestration.^49,50^ Genes from these categories (*Mt1, Mt2, Cxcl2*) are also expressed in aCap endothelial cells from PUUC lungs at 4 h, cells that do not significantly downregulate ribosomal genes. It is possible that the zinc-related effects reported to promote DC maturation, inflammatory T cell polarization, and macrophage ROS production may contribute to an antimicrobial immune function in these fibroblasts and endothelial cells at such an early timepoint.^51-53^

In the MPLA+PUUC combination specifically, the response is mainly related to proinflammatory cytokine synthesis, neutrophil chemotaxis into the lung, and ion sequestration. In the MPLA+PUUC vs. PUUC comparison, structural cells like *Ebf1*+ fibroblasts, vein endothelial cells, and adventitial fibroblasts were the strongest expressers of a triad of highly synthesized cytokines: *Cxcl2, Ccl3*, and *Ccl4*. They also express the heme scavenging protein haptoglobin (*Hp*) in MPLA+PUUC lungs. Haptoglobin is an acute phase reactant with isoforms that have been shown to upregulate MyD88-mediated inflammation,^54^ promote immunological memory to gram negative bacteria,^55^ and support B cell differentiation.^56^ Taken together, these results suggest that combination PAMPs can induce structural cells to support an antibacterial or antiviral state by cooperating with other cells such as lymphocytes and neutrophils.

Although neutrophil lung invasion was most pronounced in lungs with MPLA+PUUC NPs delivered, it is interesting that there was a modest increase in neutrophils at 24 h in lungs with PUUC NPs. Neutrophils are generally associated with bacterial infection response via their expression of bacteria-specific receptors (like TLR4) and possession of antibacterial granules.^57^ However, emerging research has illuminated the role of neutrophils in antiviral responses, noting the presence of neutrophils in BAL fluid of multiple viruses, including influenza A,^58^ respiratory syncytial viruses (RSV),^59^ and SARS-CoV-2.^60^ Lung neutrophils may contribute to the antiviral state, but they may also assist in pathological inflammation. Our data contribute to this discussion by showing that in lungs with PUUC NPs or MPLA+PUUC NPs, as early as 4 h, neutrophils exist along a defined spectrum of states characterized by the differential expression of granule proteins, cytokine signaling, and inflammatory modulators that can be computed from RNA velocity. For example, in neutrophils from the PUUC and MPLA+PUUC NP groups, granule-related transcripts *S100a8* and *S100a9* decreased while the cytokine transcripts *Il1a, Il1b*, and *Cxcl10* were increased over latent time.

It is possible that this decrease in granule proteins is a continuation of the neutrophil maturation process. These immature neutrophils are likely a part of the “left shift” of relatively immature cells fast-tracked out of the bone marrow during early acute inflammation.^61^ Given that neutrophils toward the beginning of latent time score high in established early markers like *Camp, Ngp*, and *Mmp8* and those toward the end of latent time tend to score higher in interferon-responsive genes, our results support a model in which interferons enforce an antiviral state in neutrophils as they enter the lung tissue.^39^

Our data also suggest that early neutrophil chemotaxis is under positive regulation, as neutrophils in the mid-to-late stages of latent time express themselves express neutrophil chemoattractants like *Cxcl10*.^62^ Combined with the expression of *Cxcl10* and other C-X-C chemokines by most other cell types, especially by fibroblasts and endothelial cells in the lungs with MPLA+PUUC NPs, the overall ubiquity of proinflammatory cytokine and chemokine synthesis explain the increase in neutrophils at 24 h.

Understanding the full spectrum of lung neutrophil activity may be limited by the present study. There is some early evidence of pro-apoptotic transcripts in the late-stage N0 neutrophil cluster with the expression of *Ninj1*, which regulates plasma membrane rupture during neutrophil death.^63^ However, there was little overall enrichment in canonical transcripts or gene categories for apoptosis or necroptosis in any neutrophil subcluster. Given that apoptosis is commonly the ultimate lifecycle end of neutrophils,^64,65^ it is possible that further time points past 24 h could yield valuable information about how neutrophil progression contributes to antimicrobial responses and lung pathology.

With or without MPLA, we found that RIG-I stimulation is associated with a general decrease in transcripts related to translation, particularly those encoding ribosomal proteins. This finding seems to fit with other antiviral regulatory mechanisms that function to inhibit viral protein synthesis including RNAse-L-mediated degradation of cellular mRNA (especially ribosomal protein mRNA),^66^ IFIT1-mediated inhibition of the 43S preinitiation complex (among other translational components), or zinc finger antiviral protein (ZAP) inhibition of transcripts containing certain CpG elements.^20,67^ Because these defensive anti-translational mechanisms are all under at least partial control of interferons, it is not surprising that we find transcriptional downregulation of ribosomal protein genes in a lung that is also showing strong evidence of response to interferons. This information correlates with experimental evidence that PEGylated IFN-a/ribavirin produces an analogous decrease in ribosomal protein transcripts in PBMCs from HCV patients.^68^ Interestingly, a similar broad decrease in ribosomal protein transcripts was found in the late convalescent stage of COVID-19.^69^ Based on these correlates, it is plausible that early interferons result in a transient transcriptional decrease in ribosomal biogenesis as an antiviral defense. It is unclear by what mechanism this decrease is accomplished. In eukaryotes, mTOR is known to ultimately drive synthesis of rRNA and ribosomal proteins to support cell growth.^70^ However, there is little evidence of a decrease in mTOR-related signaling pathways in the present data. It is possible that other signaling pathways downstream of IFNs are responsible for this effect especially considering the ubiquity of ISG expression in our data. It should be noted that rRNA is not expanded in the PCR amplification used our methods and that while ribosomal proteins are synthesized by RNA pol III, rRNA is synthesized by RNA pol I and is under different regulatory control.

Only NK cells from both treatment groups show no significant downregulation of ribosomal function in response to both treatments at 4 h. Indeed, either NK cell populations from PUUC or MPLA+PUUC lungs showed similar gene expression profiles. These cells show upregulation of NK-kB inhibitors (*Nfkbia;* protein IκBα) and downregulation of the Natural Cytotoxicity Triggering Receptor-1 transcript (*Ncr1*; protein NKp48 or NCR1). These results may represent an initial signaling through these pathways that is undergoing autoregulation via transcriptional control. NF-kB signaling is well-understood to be regulated by negative feedback by synthesis of pathway inhibitors like IκBα,^71,72^ and NK cell activation is similarly associated with a decrease in NCR1 expression on the cell surface,^73,74^ in addition to other stimulatory receptors like CD16 which share similar signaling pathways.^75-77^ These NK cells also demonstrate concomitant upregulation of genes in the P38-MAPK pathway, particularly *Gadd45b* and *Per1*. Both genes were noted by several recent scRNA-seq studies to be associated with the novel mNK_Sp3 / transNK3 population of splenic macrophages which may represent a heterogenous transient activated NK state composed of NK cells from multiple developmental stages.^78,79^ NK cells with this transcriptional phenotype (*Pim1, Nfkbia, Gadd45b*) were also reported to be enriched in IL-33 treated mice and were termed “signaling/inflammatory-chemokine-expressing NK cells” in a related scRNA-seq study.^80^ Although P38-MAPK signaling is typically associated with cellular proliferation, differentiation, and cell death,^81^ it is also involved in NK cell activation,^82^ including TLR3-mediated cytokine signaling in response to the PAMP poly(I:C).^83^ Given that this change is observed in our data with PAMP treatment via PUUC +/-MPLA, our results further support the concept of *Nfkbia, Pim1*, and *Gadd45b* as markers of a particular form of NK cell activation. In the scRNA-seq studies mentioned above,^78-80^ it is important to note that, like our study, this activation was observed is in the absence of foreign protein and may not necessarily indicate the ADCC function of NK cells. Rather, NK cells at this early stage (4 h) may be maintaining their ribosome function to support later cytokine production. This conclusion is supported by the differential regulation of actin filament-related GSEA categories in NK cells at this timepoint which may indicate rearrangement of cytoskeletal elements for secretory function.

Overall, it should be noted that our data suggest that cells can take on broad yet overlapping immune function in the presence of multiple external signals. Whereas PUUC +/-MPLA is associated with a ubiquitous expression of ISGs in many different cell types at 24 h, the addition of MPLA to PUUC additionally supports antibacterial function by way of the synthesis of additional antibacterial transcripts. For example, compared to AMs from murine lungs administered PUUC NPs, AMs in lungs with MPLA+PUUC NPs preferentially express transcripts for the ion sequestering regulator hepcidin (*Hamp*), ferritin heavy subunit 1 (*Fth1*), and the antibacterial siderophore lipocalin 2 (*Lcn2*). In the present data, *Hamp* is only significantly expressed in AMs in lungs with MPLA+PUUC NPs at 4 h. Considering previous reports that neither IFN-γ, TNF-a, IL-1, or IL-6 alone are responsible for *Hamp* expression in AMs at 4 h and that such expression has been reported with LPS stimulation alone,^22^ it is possible that these iron sequestering responses are in-part related to direct MPLA activation of AM-expressed TLR4. It seems that this expression fits into a general scheme of antibacterial ion binding in lungs with MPLA+PUUC NPs. Not only does iron appear to be a target of these functions, as evidenced by the expression of *Lcn2* in endothelial cells, epithelial cells, fibroblasts, lymphocytes, and monocytes, but manganese and zinc also appear as ion targets in the same cells by the expression of S100 (*S100a8, S100a9*) and metallothionein (*Mt1, Mt2*) transcripts respectively.^50,84^ These functions likely function to reduce access to metals required for bacterial growth and may doubly promote antimicrobial intracellular signaling.^85,86^

## CONCLUSION

Our results demonstrate that nanoparticulate delivery of RIG-I agonist PUUC with or without TLR4 agonist MPLA can produce an early immune response in structural, lymphoid, and myeloid cells characterized by a broad decrease in translation-related proteins within 4 hours followed by an essentially ubiquitous increase in interferon-responsive genes, including cytokines and chemokines related to neutrophil chemotaxis at 24 hours. These neutrophils appear to further increase pro-chemotactic mediators as they progress along a transcriptional trajectory starting with synthesis of classical granule proteins followed by synthesis of proinflammatory signaling molecules. With the addition of MPLA to PUUC, the general immune function of the lung is characterized by further cytokine secretion and the synthesis of antimicrobial ion-binding proteins particularly by alveolar macrophages, endothelial cells, and fibroblasts. Overall, we show that particulate delivery of PAMPs can be tuned to direct early innate inflammation in the lung. Future studies should explore how these processes could be exploited as a form of antiviral or antimicrobial therapy or prophylaxis in challenge models.

## ACKNOWLEDGEMENTS

We thank Pallab Pradhan for his advice and consultation during the early data analysis. We acknowledge the use of the Engineered Biosystems Building Physiological Research Lab for animal experiments and Biopolymer Characterization Core for the preparation of nanoparticles. The particle synthesis schematic and analytical overview figure was made with BioRender.com. We acknowledge funding from the Georgia Tech Foundation, NIH NIAID U01 AI124270, and the Robert A. Milton Chaired Professorship to Krishnendu Roy.

