## Supplemental Figures for "Single-Cell Epitope-Transcriptomics Reveal Lung Stromal and Immune Cell Response Kinetics to Nanoparticle-delivered RIG-I and TLR4 Agonists"

A

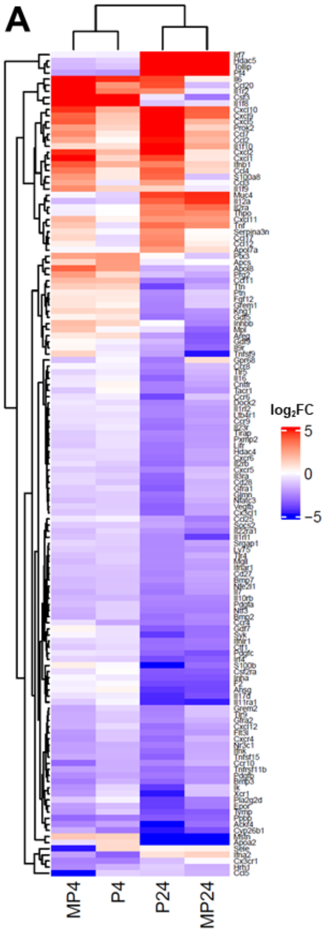


B

#### Functional Enrichment Analysis

| Treatment | Category | Term | Fold Enrichment | FDR |
|---|---|---|---|---|
| P4 | GOTERM_CC | GO:0005615~extracellular space | 1.40 | 3.87E-02 |
| | GOTERM_CC | GO:0005576~extracellular region | 1.35 | 7.31E-02 |
| | UP_KEYWORDS | Secreted | 1.35 | 2.77E-01 |
| | INTERPRO | IPR018048:CXC chemokine, conserved site | 4.54 | 4.73E-01 |
| | INTERPRO | IPR001089:CXC chemokine | 4.09 | 4.73E-01 |
| MP4 | UP_KEYWORDS | Cytokine | 1.66 | 5.13E-05 |
| | GOTERM_MF | GO:0005125~cytokine activity | 1.57 | 6.58E-04 |
| | SMART | Intercrine alpha family (small cytokine C-X-C) | 2.27 | 3.53E-03 |
| | UP_KEYWORDS | Chemotaxis | 2.05 | 1.47E-02 |
| | INTERPRO | IPR001811:Chemokine interleukin-8-like domain | 2.35 | 2.12E-02 |
| P24 | SMART | Intercrine alpha family (small cytokine C-X-C) | 3.26 | 1.45E-04 |
| | INTERPRO | IPR001811:Chemokine interleukin-8-like domain | 3.25 | 1.72E-03 |
| | GOTERM_BP | GO:0060326~cell chemotaxis | 3.10 | 3.37E-02 |
| | INTERPRO | IPR018048:CXC chemokine, conserved site | 5.40 | 1.96E-02 |
| | GOTERM_BP | GO:0030593~neutrophil chemotaxis | 2.81 | 5.21E-02 |
| | KEGG_PATHWAY | mmu04620:Toll-like receptor signaling pathway | 2.42 | 3.69E-02 |
| MP24 | INTERPRO | IPR018048:CXC chemokine, conserved site | 10.22 | 3.99E-04 |
| | INTERPRO | IPR001089:CXC chemokine | 9.20 | 4.74E-04 |
| | SMART | Intercrine alpha family (small cytokine C-X-C) | 3.73 | 2.11E-03 |
| | INTERPRO | IPR001811:Chemokine interleukin-8-like domain | 4.11 | 2.97E-03 |
| | GOTERM_MF | GO:0045236~CXCR chemokine receptor binding | 8.34 | 1.31E-02 |

C

#### P24: Toll-like Receptor Signaling Pathway

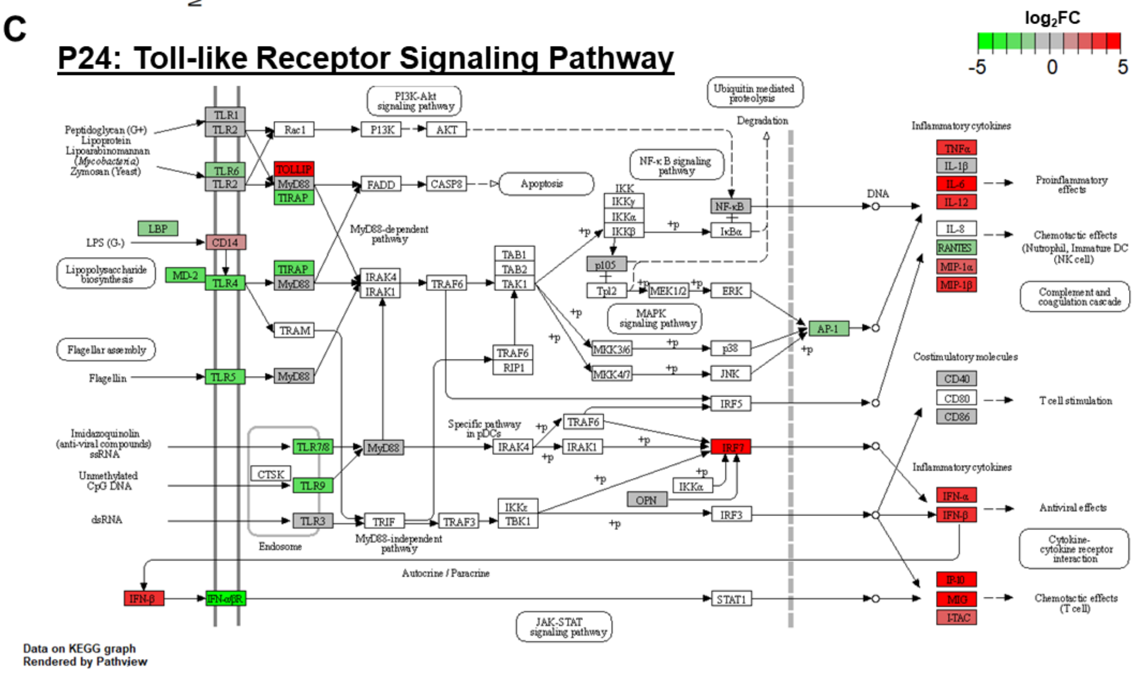

**Figure S1. PUUC and MPLA+PUUC stimulate proinflammatory cytokine and chemokine secretion in the mouse lung.** Mice were intranasally administered saline or nanoparticles (4 mg/mouse) with PUUC or MPLA+PUUC for 4 or 24 hours. Doses of 20 ug PUUC and 24 ug MPLA were used per mouse. Lungs were processed into single cell suspensions, and RNA was isolated and purified. cDNA was synthesized and pooled from three mice per treatment group into a 384-well $RT^2$ Profiler PCR Arrays using genes from the Mouse Inflammatory Response & Autoimmunity (Qiagen) set. Gene expression for treatment groups was normalized to control groups. **A)** All genes with at least one treatment group showing a $\log_2$-fold change greater or less than 2.5 were shown in a heatmap arranged with hierarchical clustering. Heatmap is colored with a $\log_2$FC cutoff of ±5. **B)** Upregulated genes with at least a fold change greater than 1.5 were inputted into the Database for Annotation, Visualization, and Integrated Discovery (DAVID) using the entire 375 microarray gene set as background with DAVID-defined defaults. Select functional categories are shown from Gene Ontology (GOTERM), Uniprot (UP_KEYWORDS), INTERPRO, SMART, and KEGG databases. **C)** Due to functional enrichment in this area, full microarray data for the PUUC 24 h group is depicted on the KEGG toll-like receptor signaling pathway (mmu0460) using a cutoff of $\log_2$FC cutoff of ±5. Genes not included in the microarray data are not shaded in the pathway. Heatmap generated with ComplexHeatmap package in R. KEGG graph rendered by Pathview in R v 4.1.0.

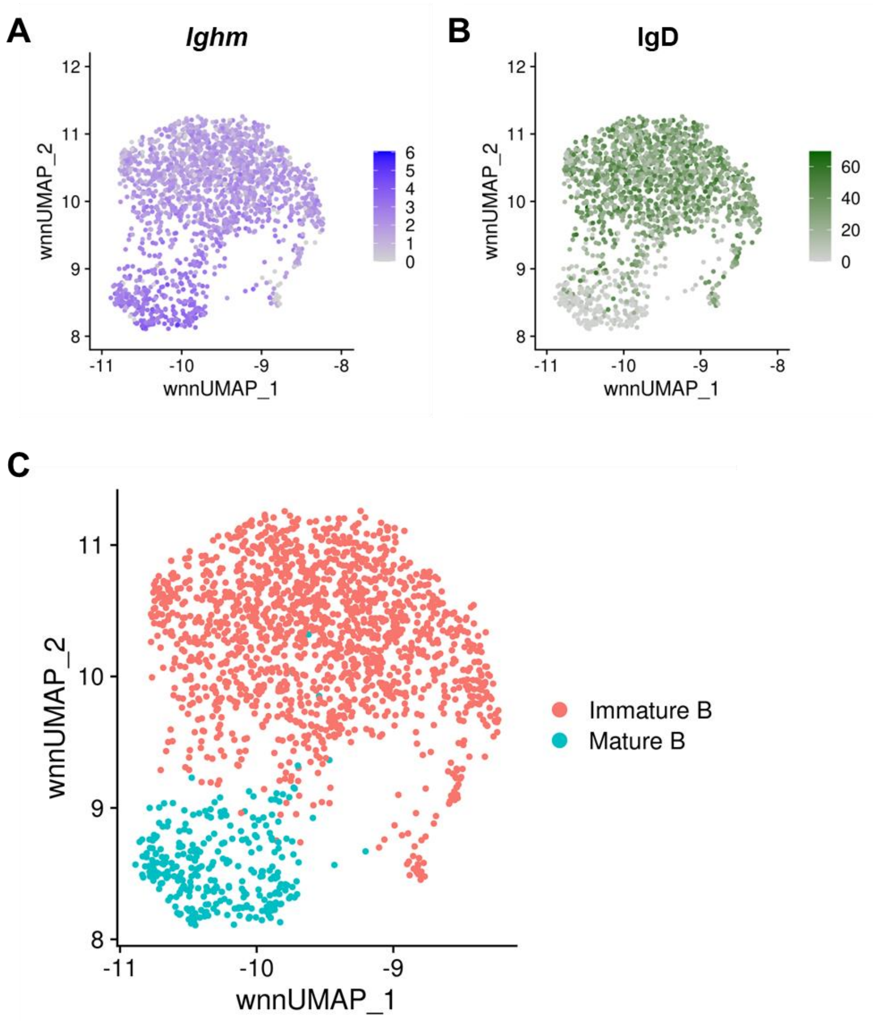


**Figure S2. B cell subsets defined by transcript and surface marker expression. A)** *Ighm* transcript expression was visualized over the B cell subset from the larger data set. **B)** Surface IgD ADT expression was likewise visualized. **C)** The final cell identities based on WNN multimodal clustering are shown for reference. All figures were generated in R 4.1.0 with Seurat V4.

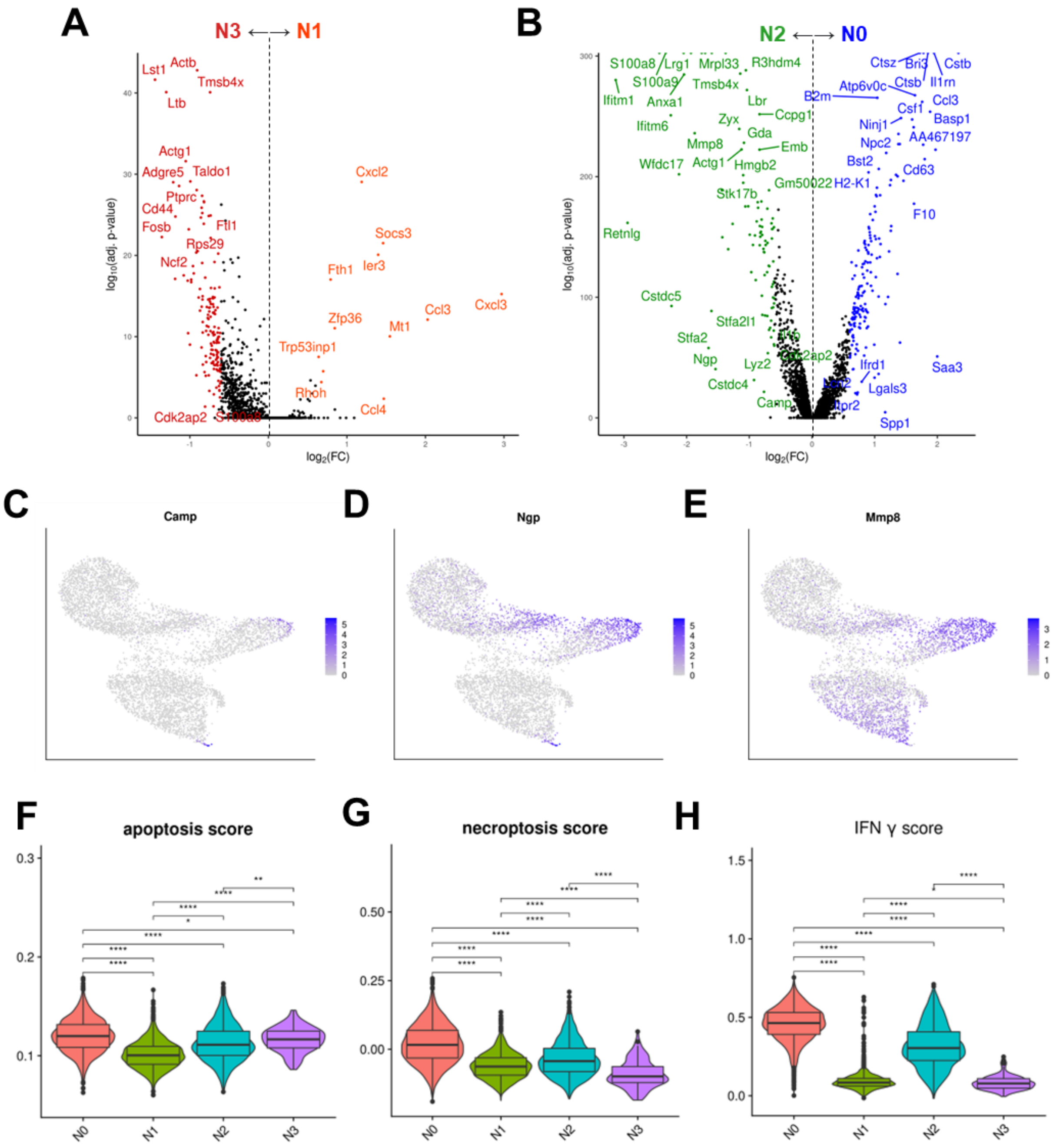


**Figure S3. Neutrophil subcluster characteristics.** Differentially expressed genes (DEGs) between neutrophil clusters **A)** N1 vs. N3 and **B)** N0 vs. N2 were visualized with volcano plots. Neutrophil gene expression data for all treatments was visualized with UMAP without integration and expression of **C)** *Camp*, **D)** *Ngp*, and **E)** *Mmp8* were plotted. Using AddModuleScore, expression of genes related to **F)** apoptosis (positive regulation of apoptotic process, GO:0043065), **G)** necroptosis (necroptosis, GO:0070266), and **H)** interferon gamma response (HALLMARK_INTERFERON_GAMMA_RESPONSE) were computed and visualized with violin and box plots. For multiple comparisons, statistical significance was calculated with a Kruskal-Wallis test. *p ≤ 0.05, **p ≤ 0.01, ***p ≤ 0.001, ****p ≤ 0.0001 for all graphs. For volcano plots, genes were colored based on clusters in Figure 3B, and a sample of differentially expressed genes with $log_2FC > 0.5$ or $< 0.5$ and $P_{adj} < 0.05$ were labeled with ggplot2. All figures were generated in R 4.1.0 with Seurat V4.

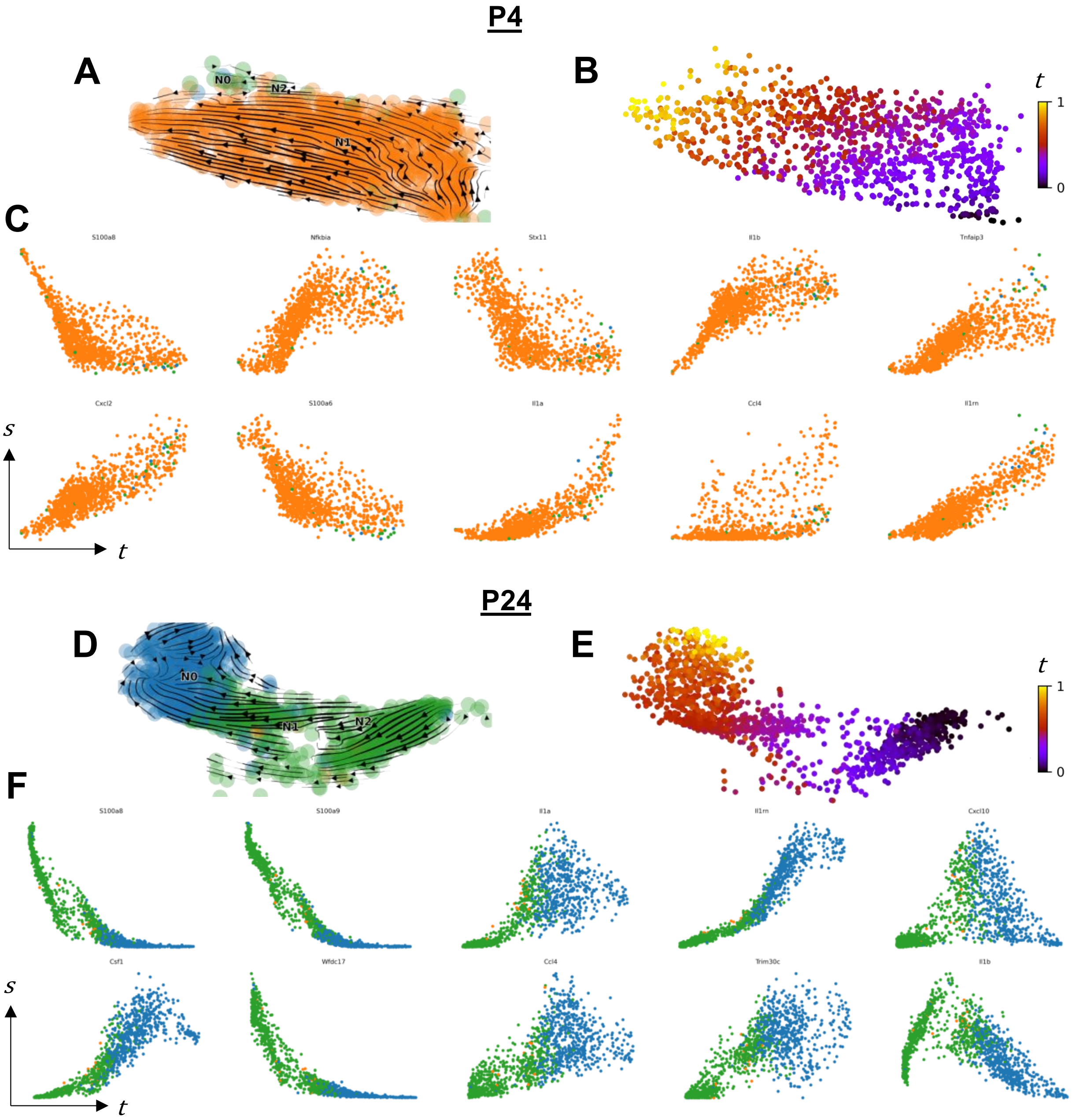


**Figure S4. Trajectory analysis of invading neutrophil gene expression along latent time from PUUC stimulated lungs at 4 and 24 hours. A)** RNA velocity profiles were computed with scVelo using dynamic modeling for neutrophils from the PUUC 4hr (P4) treatment group. RNA velocity embeddings were visualized on the original UMAP. **B)** Based on RNA velocity modeling, neutrophils were placed into a continuous latent time trajectory, *t*. **C)** Genes were selected from a group of the top 15 most-likely driver genes in the dynamic model, and expression dynamics of spliced transcripts, *s*, were visualized across latent time. Panels **D-F**) Indicate the same analysis for neutrophils from the PUUC

24h (P24) treatment group. Colors in phase portraits correspond to cluster colors in Figure 3B. UMAP coordinates, module scores, and clustering were computed with Seurat V4 in R 4.1.0. Veclocyto and scVelo were computed with Python 3.7.

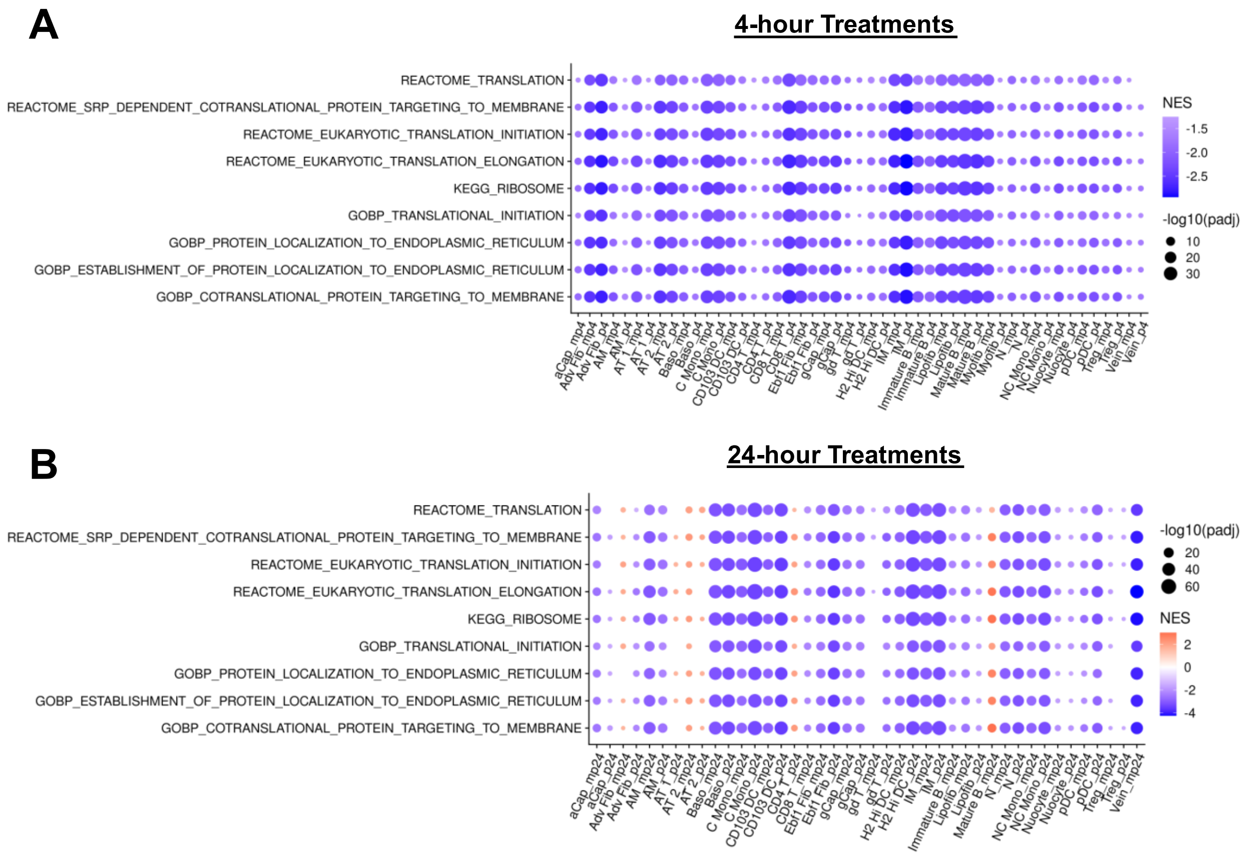


**Figure S5. Ribosome gene expression for select KEGG, REACTOME, and Gene Ontology GSEA categories for PUUC and MPLA+PUUC treated lungs at 4 and 24 hours.** Gene set enrichment analysis (GSEA) for differentially expressed genes (DEGs) were computed with FGSEA for all cell types and treatment groups compared to the naïve group. Significant GSEA results ($P_{adj} < 0.05$) for selected ribosome-related sets were visualized with dot plots at **A)** 4 h and **B)** 24 h. Gene sets for analysis included using the following categories as provided by the msigdbr package: MSigDB Hallmark gene sets, KEGG pathways, Gene Ontology Biological Processes (GOBP), and REACTOME pathways.

### NK Cell 4h DEGs

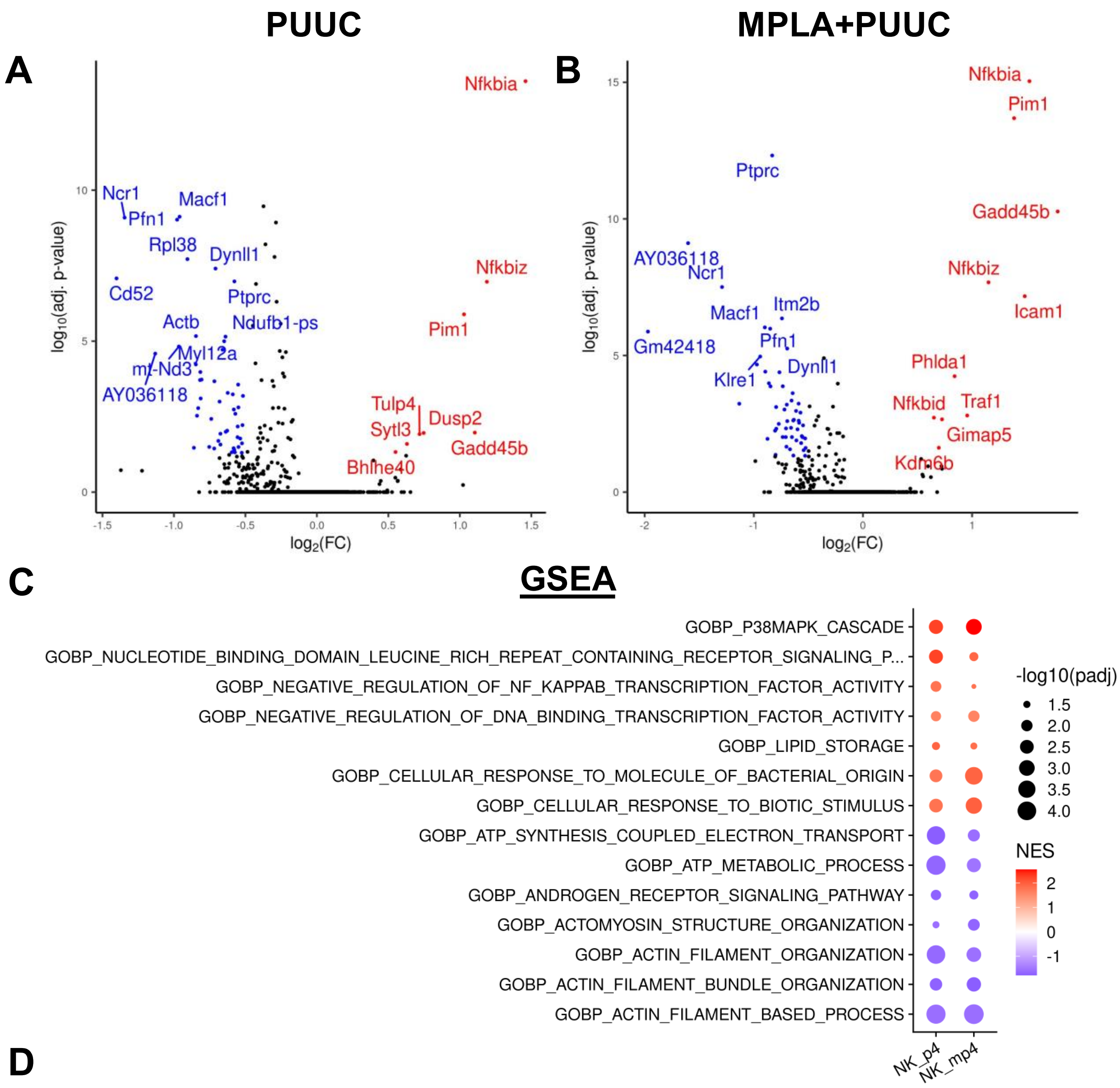


**D**

| | Pathway | adj. P | NES | Leading Edge |
|---|---|---|---|---|
| P4 | GOBP_P38MAPK_CASCADE | 2.54E-03 | 2.27 | *Gadd45b, Per1, Il1b, Zc3h12a, Dusp1, Ptpn22, Pja2, Zfp36l1, Gadd45a* |
| | GOBP_NUCLEOTIDE_BINDING_DOMAIN_LEUCINE_RICH_REPEAT... | 2.87E-03 | 2.26 | *Nfkbia, Tnfaip3, Birc3, Birc2, Rela, Ptpn22, Irak2, Ubc, Nod1, Rps27a* |
| | GOBP_POSITIVE_REGULATION_OF_CD4_ALPHA_BETA_T_CELL_ACTIVATION | 7.91E-03 | 2.19 | *Nfkbiz, Nfkbid, Ifng, Il4ra, Rara, Tgfbr2, Socs5, Myb* |
| | KEGG_CYTOSOLIC_DNA_SENSING_PATHWAY | 1.87E-02 | 2.05 | *Nfkbia, Il1b, Rela* |
| | REACTOME_EPHB_MEDIATED_FORWARD_SIGNALING | 3.12E-03 | -1.87 | *Actb, Cfl1, Actr2, Arpc1b, Cdc42, Arpc5, Actr3, Rock1, Rhoa, Fyn* |
| | GOBP_RIBONUCLEOSIDE_TRIPHOSPHATE_BIOSYNTHETIC_PROCESS | 7.58E-04 | -1.89 | *Tmsb4x, Atp5e, Atp5k, Atp5j2, Aldoa, mt-Co2, Atp5j, Nme1, Atp5h* |
| | GOBP_LOCALIZATION_WITHIN_MEMBRANE | 3.61E-03 | -1.9 | *Itgb2, Flna, Cd2, Ptprc, Itga4, Dock2, Itgal, Itgb7, Atp1b1, Synj2bp* |
| | REACTOME_CELL_SURFACE_INTERACTIONS_AT_THE_VASCULAR_WALL | 1.10E-04 | -1.94 | *Itgb2, Cd2, Fcer1g, Dok2, Spn, Pik3r1, Itgb1, Itga4, Itgal, Itgam* |
| MP4 | GOBP_P38MAPK_CASCADE | 9.18E-04 | 2.55 | *Gadd45b, Zc3h12a, Il1b, Per1, Gadd45g, Zfp36, Zfp36l1, Ager, Gadd45a* |
| | KEGG_NOD_LIKE_RECEPTOR_SIGNALING_PATHWAY | 4.66E-03 | 2.12 | *Nfkbia, Cxcl2, Il1b, Tnfaip3, Cxcl1, Rela, Birc3, Casp4* |
| | GOBP_CELLULAR_RESPONSE_TO_BIOTIC_STIMULUS | 6.07E-04 | 1.99 | *Nfkbia, Icam1, Cxcl2, Ccl3, Zc3h12a, Il1b, Tnfaip3, Cxcl1, Zfp36, Rela* |
| | GOBP_POSITIVE_REGULATION_OF_CD4_ALPHA_BETA_T_CELL_ACTIVATION | 1.24E-02 | 1.99 | *Nfkbiz, Nfkbid, Ifng, Socs5, Il4ra, Rara* |
| | GOBP_REGULATION_OF_MYELOID_LEUKOCYTE_MEDIATED_IMMUNITY | 2.67E-03 | -1.85 | *Itgb2, H2-T23, Rac2, Adgre5, Ccr2, Cd84, Itgam, Tyrobp, Vamp8, Ddx21* |
| | GOBP_INTEGRIN_MEDIATED_SIGNALING_PATHWAY | 9.18E-04 | -1.87 | *Itgb2, Itgb1, Txk, Cdc42, Flna, Zyx, Plek, Itgal, Itgam, Itga2, Itga4, Nedd9* |
| | GOBP_CELLULAR_DEFENSE_RESPONSE | 2.20E-03 | -1.87 | *Ncr1, Spn, Itgb1, Klrc1, Ccr5, Prf1, Ccr2, Klrc2, Tyrobp, Klrg1* |
| | KEGG_FOCAL_ADHESION | 2.15E-05 | -1.92 | *Diaph1, Pik3r1, Actb, Itgb1, Cdc42, Flna, Zyx, Myl12a, Rock1, Bcl2* |

**Figure S6. Natural Killer cells demonstrate a similar NF-kB-related gene-expression profile at 4 hours following PUUC or MPLA+PUUC stimulation.** Differentially expressed genes (DEGs) between NK cells for the **A)** PUUC vs. naïve and **B)** MPLA+PUUC vs. naïve groups at 4 h were visualized with volcano plots. **C)** GSEA was computed from DEGs, and the top 7 most significant up- and down-regulated gene sets were visualized on a dot plot. **D)** Four of the top 5 enriched up- and down-regulated GSEA categories are depicted in a table with genes from the leading edge shown. For volcano plots, a sample of differentially expressed genes with $log_2FC > 0.5$ or $< 0.5$ and $P_{adj} < 0.05$ were labeled with ggplot2. All figures were generated in R 4.1.0

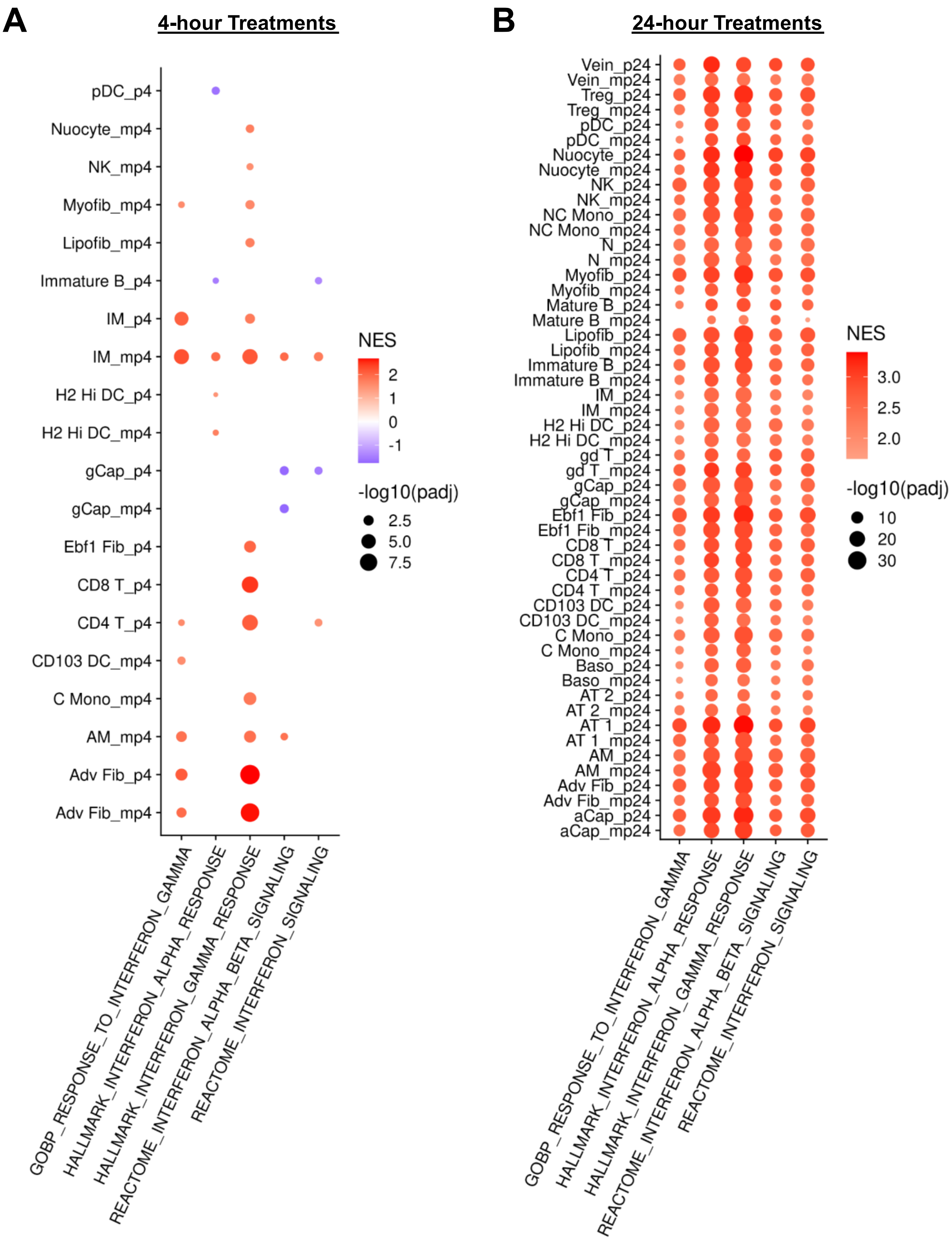
A
4-hour Treatments
pDC_p4
Nuocyte_mp4
NK_mp4
Myofib_mp4
Lipofib_mp4
Immature B_p4
IM_p4
IM_mp4
H2 Hi DC_p4
H2 Hi DC_mp4
gCap_p4
gCap_mp4
Ebf1 Fib_p4
CD8 T_p4
CD4 T_p4
CD103 DC_mp4
C Mono_mp4
AM_mp4
Adv Fib_p4
Adv Fib_mp4
GOBP_RESPONSE_TO_INTERFERON_GAMMA
HALLMARK_INTERFERON_ALPHA_RESPONSE
HALLMARK_INTERFERON_GAMMA_RESPONSE
REACTOME_INTERFERON_ALPHA_BETA_SIGNALING
REACTOME_INTERFERON_SIGNALING
NES
2
1
0
-1
-log10(padj)
2.5
5.0
7.5
B
24-hour Treatments
Vein_p24
Vein_mp24
Treg_p24
Treg_mp24
pDC_p24
pDC_mp24
Nuocyte_p24
Nuocyte_mp24
NK_p24
NK_mp24
NC Mono_p24
NC Mono_mp24
N_p24
N_mp24
Myofib_p24
Myofib_mp24
Mature B_p24
Mature B_mp24
Lipofib_p24
Lipofib_mp24
Immature B_p24
Immature B_mp24
IM_p24
IM_mp24
H2 Hi DC_p24
H2 Hi DC_mp24
gd T_p24
gd T_mp24
gCap_p24
gCap_mp24
Ebf1 Fib_p24
Ebf1 Fib_mp24
CD8 T_p24
CD8 T_mp24
CD4 T_p24
CD4 T_mp24
CD103 DC_p24
CD103 DC_mp24
C Mono_p24
C Mono_mp24
Baso_p24
Baso_mp24
AT 2_p24
AT 2_mp24
AT 1_p24
AT 1_mp24
AM_p24
AM_mp24
Adv Fib_p24
Adv Fib_mp24
aCap_p24
aCap_mp24
GOBP_RESPONSE_TO_INTERFERON_GAMMA
HALLMARK_INTERFERON_ALPHA_RESPONSE
HALLMARK_INTERFERON_GAMMA_RESPONSE
REACTOME_INTERFERON_ALPHA_BETA_SIGNALING
REACTOME_INTERFERON_SIGNALING
NES
3.0
2.5
2.0
-log10(padj)
10
20
30

**Figure S7. Following PUUC and MPLA + PUUC treatment, GSEA of differentially expressed genes reveals mild upregulation of interferon-related gene sets at 4 hours followed by large increases in diverse cell types at 24 hours.** Gene set enrichment analysis (GSEA) for differentially expressed genes (DEGs) were computed with FGSEA for all cell types and treatment groups compared to the naïve group. Significant GSEA results ($P_{adj}$ < 0.05) for selected interferon-related sets were visualized with dot plots at **A)** 4 h and **B)** 24 h. Gene sets for analysis included using the following categories as provided by the msigdbr package: MSigDB Hallmark gene sets, KEGG pathways, Gene Ontology Biological Processes (GOBP), and REACTOME pathways.

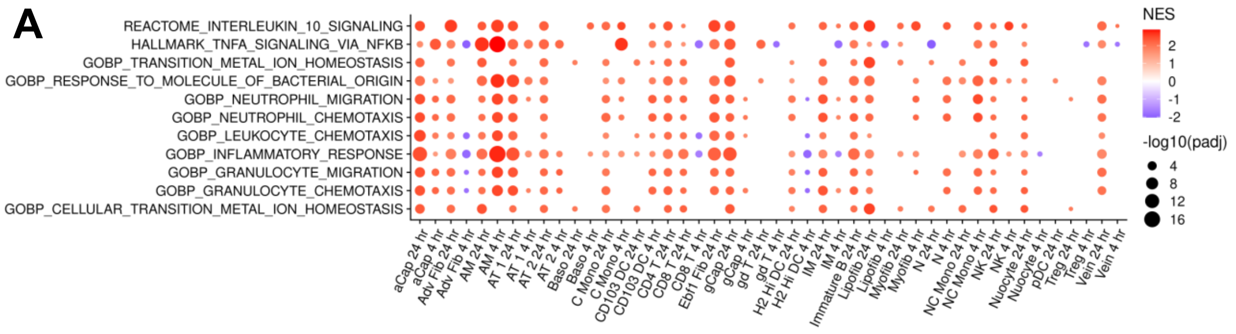


**Figure S8. GSEA of diverse cell types in the MPLA+PUUC vs. PUUC stimulated lung reveals enrichment in genes associated with neutrophil chemotaxis, transition metal ion homeostasis, and TNF$\alpha$-related signaling.** Gene set enrichment analysis (GSEA) for differentially expressed genes (DEGs) were computed with FGSEA for all cell types for the MP vs. P treatment groups across 4 and 24 h timepoints. **A)** Significant GSEA results ($P_{adj} < 0.05$) for selected gene sets were visualized with a dot plot. NES = Normalized enrichment score.
